# Scavenger receptor A1 participates in the phagocytosis of *Leptospira interrogans* and leads to subsequent high inflammatory responses and bacterial dissemination in leptospirosis

**DOI:** 10.1101/2020.11.27.401083

**Authors:** Yanchun Wang, Xia Fan, Lin Du, Boyu Liu, Haihan Xiao, Yan Zhang, Yunqiang Wu, Fuli Liu, Yung-Fu Chang, Xiaokui Guo, Ping He

## Abstract

Leptospirosis, caused by pathogenic *Leptospira* species, has emerged as a widespread zoonotic disease worldwide. Macrophages mediate the elimination of pathogens through phagocytosis and cytokine production. Scavenger receptor A1 (SR-A1), one of the critical receptors mediating this process, plays a complicated role in innate immunity. However, the role of SR-A1 in the immune response against pathogenic *Leptospira* invasion is unknown. In the present study, we found that SR-A1 is an important nonopsonic phagocytic receptor on murine macrophages for *Leptospira*. We also found that leptospiral LPS is the ligand of SR-A1. However, intraperitoneal injection of leptospires into WT mice presented with more severe jaundice, subcutaneous hemorrhaging, and higher bacteria burdens in blood and tissues than that of SR-A1^-/-^ mice. Exacerbated cytokine and inflammatory mediator levels were also observed in WT mice and higher recruited macrophages in the liver than those of SR-A1^-/-^ mice. Our findings collectively reveal that although beneficial in the uptake of *Leptospira* by macrophage, SR-A1 might be exploited by *Leptospira* to promote bacterial dissemination and modulate inflammatory activation, which causes a more severe infection in the host. These results provide our new insights into the innate immune response during early infection by *L. interrogans*.

## Introduction

Leptospirosis, known as Weil’s disease, is caused by pathogenic species of the genus *Leptospira*, which has emerged as the most widespread zoonotic disease worldwide (Evangelista & Coburn, 2010; Soo, Khan, & Siddiqui, 2020). Human leptospirosis is an acute febrile illness with a wide range of clinical features from mild flu-like symptoms to severe leptospirosis characterized by jaundice, bleeding, pulmonary hemorrhage, renal failure and death (Palaniappan, Ramanujam, & Chang, 2007).

Innate immune responses constitute the first line of defense against *Leptospira*. At the early stages of *Leptospira* infection, macrophages play a complicated role against *Leptospira* by phagocytosis and induction of signaling pathways to produce pro-inflammatory cytokines and antigen presentation (Ignacio Santecchia, Ferrer, Vieira, Gómez, & Werts, 2020; I. Santecchia et al., 2019; Werts, 2018).

The process of bacterial recognition and phagocytosis by macrophages has been intensively studied in recent decades (Kubelkova & Macela, 2019; Pluddemann, Mukhopadhyay, & Gordon, 2006). A comprehensive series of surface receptors have been identified as being involved in the phagocytosis of microorganisms, including integrins, Fc gamma receptors (FcγRs), mannose receptor, and the scavenger receptors, etc. (S. A. Freeman & Grinstein, 2014). In the case of *Leptospira*, our understanding of the receptors involved in phagocytosis is limited. Two phagocytic receptors described for *Leptospira* are the third complement receptor (CR3) and β2 integrin, and much less is known whether other receptors are involved in leptospiral phagocytosis (Cinco, Cini, Perticarari, & Presani, 2002; Cosate, Siqueira, de Souza, Vasconcellos, & Nascimento, 2016).

Scavenger receptor A1 (SR-A1), also called macrophage scavenger receptor (MSR) or CD204, belongs to a class of pattern recognition receptors (PRRs) expressed primarily on macrophages (PrabhuDas et al., 2017). SR-A1 was initially described as a receptor for modified lipoproteins involved in atherosclerosis development (Goldstein, Ho, Basu, & Brown, 1979). It has been identified as being involved in many critical biological processes, such as adhesion and phagocytosis (M. Freeman et al., 1991). Studies have shown that SR-A1, as a nonopsonic phagocytic receptor, can bind and phagocytose various bacteria, such as *Escherichia coli, Neisseria meningitidis, Streptococcus pneumoniae, Staphylococcus aureus* and *Listeria monocytogenes* (Arredouani et al., 2006; Dunne, Resnick, Greenberg, Krieger, & Joiner, 1994; Peiser et al., 2002; Peiser, Gough, Kodama, & Gordon, 2000). In addition, there is evidence that SR-A1 is involved in regulating innate immune responses and proinflammatory cytokine responses to pathogen infection (Areschoug & Gordon, 2009). However, no studies have been reported about the interactions of SR-A1 with *Leptospira*. Thus, we sought to evaluate the role of SR-A1 during the *Leptospira* infection.

Animal models represent essential tools in research on the pathogenic mechanism of leptospirosis. Guinea pigs and hamsters have been the most commonly used animal models for *Leptospira* infection (Gomes-Solecki, Santecchia, & Werts, 2017). We recently developed a murine model of acute and self-resolving leptospirosis by infecting adult, immunocompetent C57BL/6 mice with *L. interrogans* serovar Autumnalis strain 56606v (Xia et al., 2017). This murine leptospirosis model closely recapitulates natural disease in humans, with characteristic manifestations including prominent jaundice and pulmonary hemorrhage (Xia et al., 2017). Thus, we use WT and SR-A1^-/-^ mice to investigate the host-pathogen interactions of leptospirosis.

Our results showed that SR-A1 is an important nonopsonic phagocytic receptor for *Leptospira* ingestion. Additionally, SR-A1 contributes to regulating proinflammatory cytokine responses to *Leptospira* infection and affects the disease outcome. These results provide our new insights into the innate immune response during early infection by *L. interrogans*.

## Materials and methods

### Bacterial strains

Pathogenic *L. interrogans* serovar Autumnalis strain 56606v was kindly provided by the Institute for Infectious Disease Control and Prevention (Beijing, China), and serial passages in guinea pigs to maintain bacterial virulence. Leptospires *in vitro* were cultivated in EMJH medium at 28 °C to mid −log phase. After counting in a Petroff-Hauser chamber under dark-field microscopy, *L. interrogans* were suspended in PBS at a particular cell density for the experiment. The bacterial suspensions were then fixed for 1 h at 4 °C with 4% paraformaldehyde to inactivate the bacteria. These preparations were washed and resuspended in PBS for use.

### Cell culture

Murine bone marrow-derived macrophages (BMDMs) were generated from WT and SR-A1^-/-^ mice, as previously described with minor modifications (Marim, Silveira, Lima, & Zamboni, 2010). Briefly, tibiae and femurs from mice were flushed with PBS, and bone marrow cells were resuspended in RPMI 1640 (Gibco, USA) supplemented with 10% heat-inactivated FBS, 30% L929-conditioned medium, 1% HEPES and 1% Penicillin-Streptomycin (Gibco, USA). Cells were cultured in 10 cm Petri dishes (Nunc, Denmark) for 5 days at 37 °C in a humidified incubator with 5% CO_2_. Then, macrophages were obtained by scratching in cold PBS containing 0.5% EDTA, followed by centrifugation at 300×*g* for 10 min. The cells were washed again and subsequently cultured for experimental use.

Peritoneal macrophages (PMs) were harvested from WT and SR-A1^-/-^ mice after injection of 1 mL of 5% thioglycollate broth intraperitoneally, as previously described (Pavlou, Wang, Xu, & Chen, 2017). After lavaging the peritoneal cavity with 10 mL of cold RPMI 1640 followed by centrifugation at 300×*g* for 10 min, the cells were subsequently cultured in Petri dishes of 10% FBS RPMI 1640 for 2 h allowing macrophages to adhere.

Raw264.7 and HEK293T cells, originally from American Type Culture Collection, were cultured in DMEM (Corning, USA) containing 10% FBS and 1% Penicillin-Streptomycin maintained at 37 °C in a humidified 5% CO_2_ incubator.

### Plasmid construction and transfection

Full-length murine macrophage SR-A1 cDNA ORF was cloned into a pHBLV-U6-ZsGreen-Puro plasmid (Hanbio, China) for lentiviral production. The recombinant retrovirus was transfected into RAW264.7 and HEK293T cells according to the manufacturer’s instructions. Stable cells were screened with 800 μg/mL puromycin. Efficiency expression of SR-A1 was determined by Western blotting and flow cytometry (FCM) method.

### Leptospires stimulation and phagocytosis

Cells seeded at 2×10^5^ per well were incubated on slides in a 24-well plate under appropriate culture conditions. Cells were infected with *L. interrogans* strain 56606v at MOI of 10 to 100 paraformaldehyde-fixed bacteria or 100 live bacteria per cell, respectively. To synchronize the stage of infection, the plates were centrifuged at 100×*g* for 10 min and then co-incubated in a humidified 5% CO_2_ incubator for 1 h. The cells were washed extensively with pre-cooled PBS to stop phagocytosis and remove extracellular *L. interrogans*.

In phagocytosis experiments, cells were pre-incubated before leptospires infection with polyI (100 μg/mL, Sigma, USA), polyC (100 μg/mL, Sigma), SR-A1 monoclonal antibody 2F8 (30 μg/mL, AbD serotec, United Kingdom), isotype control rat IgG2b (30 μg/mL, AbD serotec), cytochalasin D (20 μM, Sigma), mannan from *Saccharomyces cerevisiae* (10 mg/mL, Sigma) or N-Acetyl-D-glucosamine (50 μM, Sigma) for 30 min in the condition without FBS. Rabbit anti-*L. interrogans* strain 56606v antibody was used as the specific primary antibody. FITC-conjugated (BD, USA) or Alexa Flour 647-conjugated anti-rabbit IgG (Abcam, USA) as a secondary antibody was used before permeabilization (1% paraformaldehyde and 0.5% Triton X-100), while TRITC-conjugated anti-rabbit IgG (ProteinTech, USA) was used after that. Nuclei were stained with 1 μM DAPI for 10 min, and slides were sealed and detected by laser using an Olympus Confocal microscopy.

Thus, phagocytic leptospires inside macrophages were only stained by one fluorescein-labeled antibody after permeabilization, while leptospires adhering outside were stained by diverse conjugated antibodies both pre-and post-permeabilization. Two hundred macrophages were counted in several HP fields, and the rates of macrophages with phagocytosed leptospires were calculated.

### Leptospiral LPS extraction and phagocytosis of LPS coated beads

Leptospiral LPS was extracted and purified according to the hot phenol-water method (Werts et al., 2001). Microbeads of 1 μm diameter were coated with LPS (50 μg/mL) of *L. interrogans* strain 56606v or 5% BSA at 37 °C for 1 h. Beads and cells were co-cultured for 1 h at the concentrations of MOI=10. The uptake of beads in HEK293T and BMDMs were detected by the confocal microscopy method.

### Experimental Animals

SR-A1-deficient (SR-A1^-/-^) mice on the C57BL/6 background were kindly provided by Prof. Qi Chen, Nanjing Medical University (Guo et al., 2016). Age (8 weeks old) and gender (female)-matched WT C57BL/6 mice obtained from Shanghai Jiaotong University School of Medicine were used as controls. Primary cells from BALB/c mice were used in the experiment of SR-A1 inhibition because the commercial antibody 2F8 did not block the SR-A1 of C57BL/6 mice (Daugherty, Whitman, Block, & Rateri, 2000). These mice were maintained under specific-pathogen-free conditions in the vivarium of the Experimental Animal Center at Shanghai Jiaotong University School of Medicine. All experiments were performed in strict accordance with the Regulations for the Administration of Affairs Concerning Experimental Animals. The Animal Ethics Review Committee of Shanghai Jiaotong University approved all animal procedures (project number A-2018-021).

### *Leptospira* infection with WT and SR-A1^-/-^ mice

WT and SR-A1^-/-^ mice were infected with 2×10^8^ leptospires in 200 μL PBS via the intraperitoneal (IP) route according to the method described (Ratet et al., 2014). Negative control mice were intraperitoneally injected with 200 μL PBS. The mice were bled and sacrificed at 1, 2, 3, and 5 days post-infection (dpi), respectively. The blood, liver and kidney of WT and SR-A1^-/-^ mice were collected for RNA extraction, histological and immunohistochemical analysis.

For the test of phagocytosis of PMs to leptospires *in vivo*, the mice were sacrificed at 2 and 24 h post-infection (hpi), respectively. PMs were harvested and cultured on slides in a 24-well plate. Adherent or phagocytized leptospires were labeled and detected by specific antibodies as the method mentioned above.

### Bacterial loads

The burden of leptospires in murine blood or tissues was analyzed by reverse transcription and real-time PCR according to the method described (Xia et al., 2017). The concentration of leptospires in the animal blood and tissues was quantified with an ABI 7500 PCR System (Applied Biosystems) using Power SYBR Green PCR Master Mix (Applied Biosystems). The concentration of the final PCR product (16S rRNA: 5’-AGC ACG TGT GTT GCC CTA GAC ATA-3’ and 5’-GTT GCC ATC ATT CAG TTG GGC ACT-3’) was calculated by 2^−ΔCt^ relative to GAPDH as the reference gene.

### Pathological and immunohistochemical studies

Tissues (liver and kidney) were collected and fixed in 4% paraformaldehyde and then embedded in paraffin. Tissue sections were stained with hematoxylin and eosin (H&E). Tissue injury was examined by light microscopy. Immunohistochemical staining, using *L. interrogans* strain 56606v-specific rabbit antiserum prepared in our lab, was performed using the EnVison system (Dako, Glostrup, Denmark).

### Serum biochemical analysis

The serum was collected from blood samples and stored at −80 °C. The concentrations of total bilirubin (TBIL), aspartate transaminase (AST) and serum creatinine (CREA) were measured by using a UniCel DxC 800 Synchron autoanalyzer (Beckman Coulter, Brea, CA, USA).

### Quantification of surface receptors, inflammatory mediator and cytokines in macrophages and tissues

WT and SR-A1^-/-^ PMs were infected by alive *L. interrogans* strain 56606v as MOI=100 for 2 h *in vitro*, and then 100 μg/mL gentamicin was used for 1 h to kill the extracellular leptospires. Culture medium was replaced and primary PMs were extracted for RNA detection (marked as 3 hpi). For protein detection, PMs were cultured for 1 d (1 dpi) and supernatants were collected.

TRIzol LS Reagent (Thermo Scientific, USA) was used for RNA extraction from primary cells stimulated by leptospires *in vitro*, and from blood or tissues infected by leptospires *in vivo*. Reverse transcription was used by a SuperScript III First-Strand Synthesis SuperMix Kit (Thermo Scientific), and cDNA was then subjected to real-time quantitative PCR (qPCR) with ABI 7500 PCR System (Applied Biosystems, USA). Primers were listed in the supplemental material (Supplementary Table 1). Real-time PCR data were calculated by 2^−ΔΔCt^ relative to GAPDH as the reference gene and cells without leptospiral stimulation as control.

The cytokine detection of supernatants was performed using ELISA kits (R&D Systems, USA) for mouse IL-1β, IL-6 and TNFα, according to the instructions provided by the supplier. Nitric oxide (NO) formation was quantified immediately via the Griess reaction.

### Statistical analysis

Statistical analysis was performed using SPSS version 20.0 (IBM Corporation, USA), and figures were generated using GraphPad Prism software version 5.0. The data were expressed as means ± SEM. The mean values obtained from the experiments were compared utilizing Student’s *t*-test, one-way ANOVA analysis with post-hoc test or multiple *t*-tests. A *P*-value of less than 0.05 was considered significant.

## Results

### SR-A1 is an important receptor for non-opsonic-phagocytosis of *L. interrogans* by murine macrophage

We assessed the regulation of candidate macrophage surface receptors in response to stimulation with *L. interrogans* 56606v in order to screen the receptors participating in the recognition of *Leptospira*. Integrins, Fc receptors, C-type lectins, and sialic acid-binding Ig-like lectins (siglecs), and several other receptors belonging to the scavenger receptor family were regulated by stimulation with *L. interrogans* (Fig. 1). Notably, only the expression of SR-A1 from these candidate receptors was found to be significantly increased, further suggesting the potential role of SR-A1 in the interaction of macrophages and *Leptospira*.

**Fig. 1.**
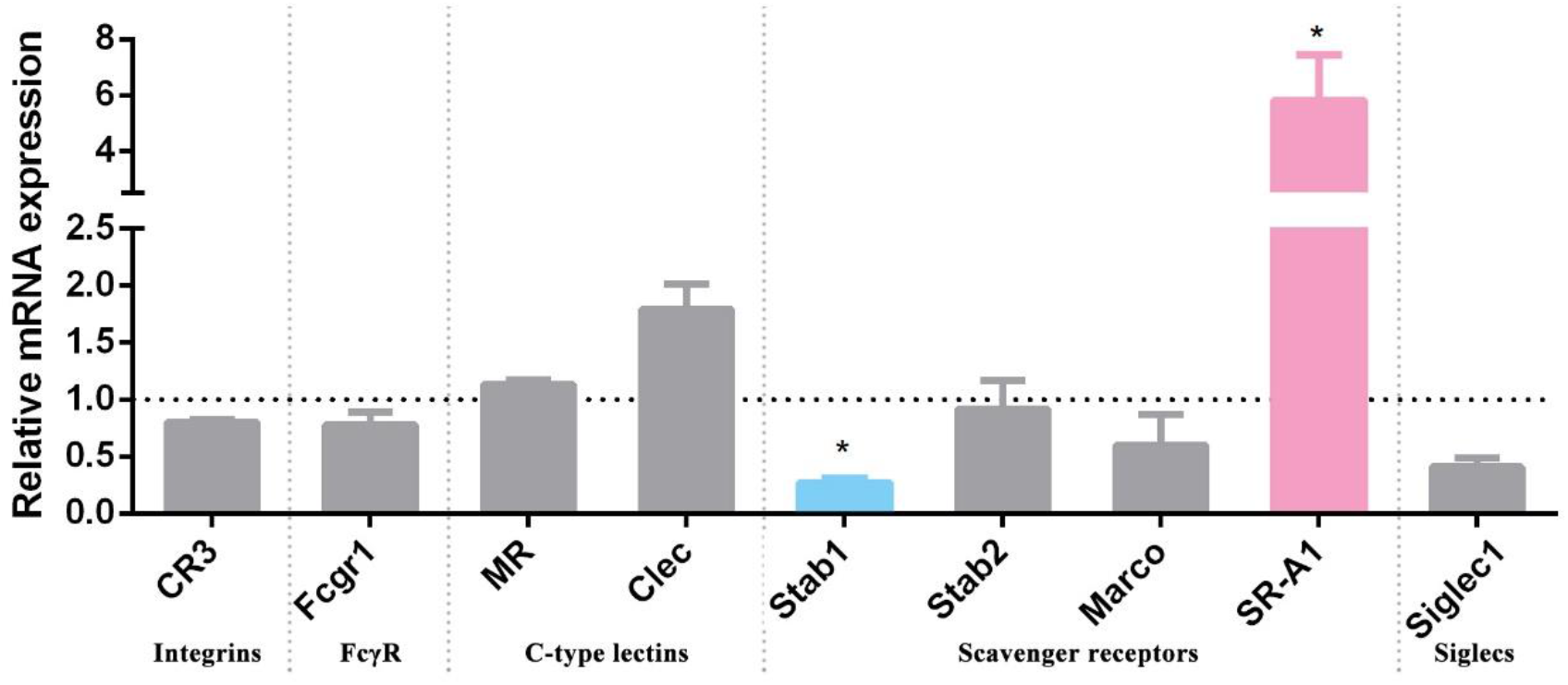
Regulation of surface receptors of PMs by stimulation with *L. interrogans* strain 56606v. PMs were incubated with *L. interrogans* strain 56606v for 3 hours. Total RNA was extracted and reverse transcribed cDNA was detected by qPCR method. Relative expression of surface receptors was calculated by 2^−ΔΔCt^ relative to GAPDH as the reference gene and cells without leptospiral stimulation as control. These data were expressed as the mean ± SEM from at least three experiments. *\*P*<*0*.*05* and significant in multiple t tests.

To determine the contribution of SR-A1 to the non-opsonic-phagocytosis of *L. interrogans* by murine BMDM, we investigated the inhibition of leptospires uptake by SR-A1 ligand polyI (a general SR inhibitor) and anti-SR-A1 2F8 (an anti-SR-A1 monoclonal antibody) without opsonins. Indirect immunofluorescence staining was used to detect the leptospires inside (red) and outside (merged in yellow for both binding of FITC-conjugated and TRITC-conjugated antibodies before and after permeabilization) of macrophages. Confocal microscopic images showed that polyI and anti-SR-A1 could significantly inhibit the phagocytosis of inactive *L. interrogans* by BMDM cells compared to the corresponding controls polyC and rIgG2b (Fig. 2). Similar results were found in inhibition experiments of phagocytosis of active *L. interrogans* by BMDM cells (Supplementary Fig. 1). The FCM method was employed further to confirm our findings (Supplementary Fig. 2). The results showed that inhibitors of SR-A1 could significantly reduce phagocytosis rates of *Leptospira* by BMDM cells. However, inhibition of the mannan receptor which was not significantly regulated by leptospiral stimulation, exhibited no inhibition to phagocytosis in *L. interrogans* strain 56606v, indicating this receptor did not participate in this process (Supplementary Fig. 3).

**Fig. 2.**
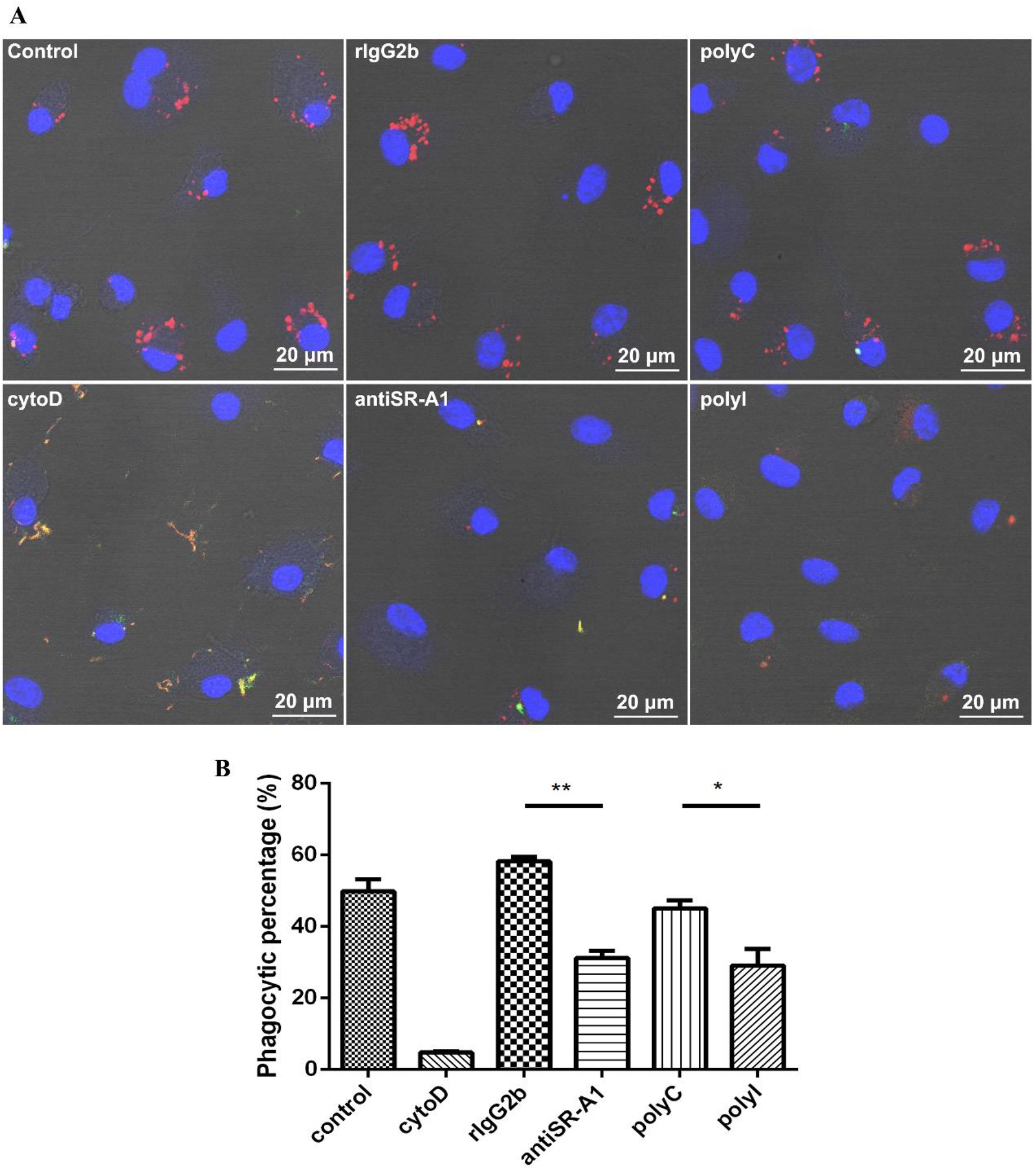
PolyI and SR-A1 monoclonal antibody (anti-SR-A1) exhibited the inhibition to the phagocytosis of *L. interrogans* strain 56606v by mouse BMDMs. BMDMs were incubated with inactive *L. interrogans* strain 56606v in FBS-free medium absence or presence of cytoD (20μM), polyI (100μg/mL) and anti-SR-A1 (30μg/mL). Corresponding concentrations of polyC and rat IgG2b (rIgG2b) isotypes were added as controls. Rabbit anti-*L. interrogans* strain 56606v was treated as a specific primary antibody, while FITC-conjugated or TRITC-conjugated anti-rabbit IgG as a secondary antibody were used before and after permeabilization. Confocal microscopic images showed leptospires inside (red) or outside (yellow) of BMDMs **(A)**. Phagocytic percentages of BMDMs phagocytizing *L. interrogans* were calculated and statistically analyzed by variance **(B)**. These data were expressed as the mean ± SEM from at least three experiments. *\*P*<*0*.*05, **P*<*0*.*01*.

To further verify the role of SR-A1 in *L. interrogans* adhesion and phagocytosis, HEK293T or RAW264.7 cells transfected with SR-A1 vector were constructed. Cell surface overexpression of SR-A1 was verified by the FCM method (Supplementary Fig. 4). SR-A1-transfected HEK293T cells showed significantly increased adhesion of inactive *L. interrogans* compared to mock cells, with a large amount of fluorescent merged leptospires visible attaching to the cell membrane (Fig. 3A). Overexpression of SR-A1 in RAW264.7 cells also showed higher phagocytic rates than mock cells (Fig. 3B). These results confirmed the SR-A1 function in adhesion and phagocytosis of leptospires.

**Fig. 3.**
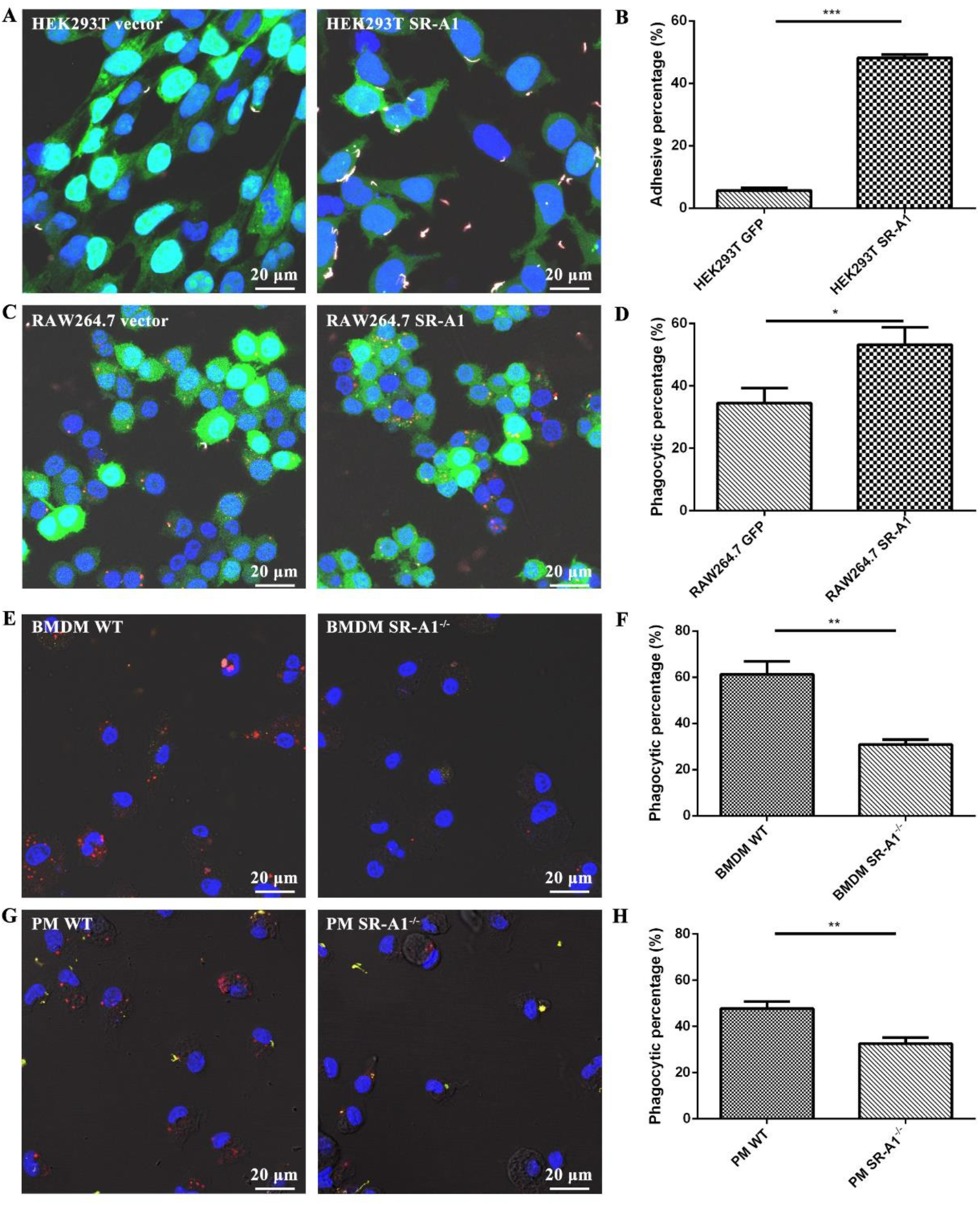
SR-A1 enhanced *L. interrogans* adhesion and phagocytosis. Lentivirus transfected HEK293T vector cells and HEK293T SR-A1 cells **(A)**, RAW264.7 vector cells and RAW264.7 SR-A1 cells **(C)** with GFP expression, WT and SR-A1^-/-^ BMDMs **(E)** or PMs **(G)** were incubated with *L. interrogans* strain 56606v at MOI=50 for 1 h in medium without FBS, respectively. Rabbit anti-*L. interrogans* strain 56606v was used as a specific primary antibody, while Alexa Fluor 647-conjugated or TRITC-conjugated anti-rabbit IgG as a secondary antibody were used before and after permeabilization, respectively. Confocal microscopic images showed leptospires inside (red) or outside (white/yellow) of cells. Positive rates of *Leptospira* adhesive cells or phagocytic cells were calculated, respectively **(B, D, F, H)**. These data were expressed as the mean ± SEM from at least three experiments. *\*P*<*0*.*05, **P*<*0*.*01, ***P*<*0*.*001*.

SR-A1 gene knockout mice were used to further characterize the role of SR-A1 in the phagocytosis of leptospires. SR-A1^-/-^ BMDMs phagocytosed about 50% less *L. interrogans* than WT BMDMs did (Fig. 3E and F). Consistent results were also achieved in PMs from SR-A1^-/-^ and WT mice (Fig. 3G and H). These data indicated that SR-A1 is an important murine macrophage receptor for non-opsonic-phagocytosis of *L. interrogans*.

### SR-A1^-/-^ macrophages were deficient in phagocytosis of *L. interrogans in vivo*

To evaluate the phagocytosis ability of SR-A1 in the presence of opsonin *in vivo*, we inoculated leptospires in the peritoneal cavity of WT and SR-A1^-/-^ mice. PMs were harvested at 2 and 24 hpi, and leptospires were stained and detected by confocal microscopy. The result showed that WT PMs have higher phagocytic rates than SR-A1^-/-^ PMs (Fig. 4). Our results showed that SR-A1 also plays a significant role in the phagocytosis of *L. interrogans in vivo*.

**Fig. 4.**
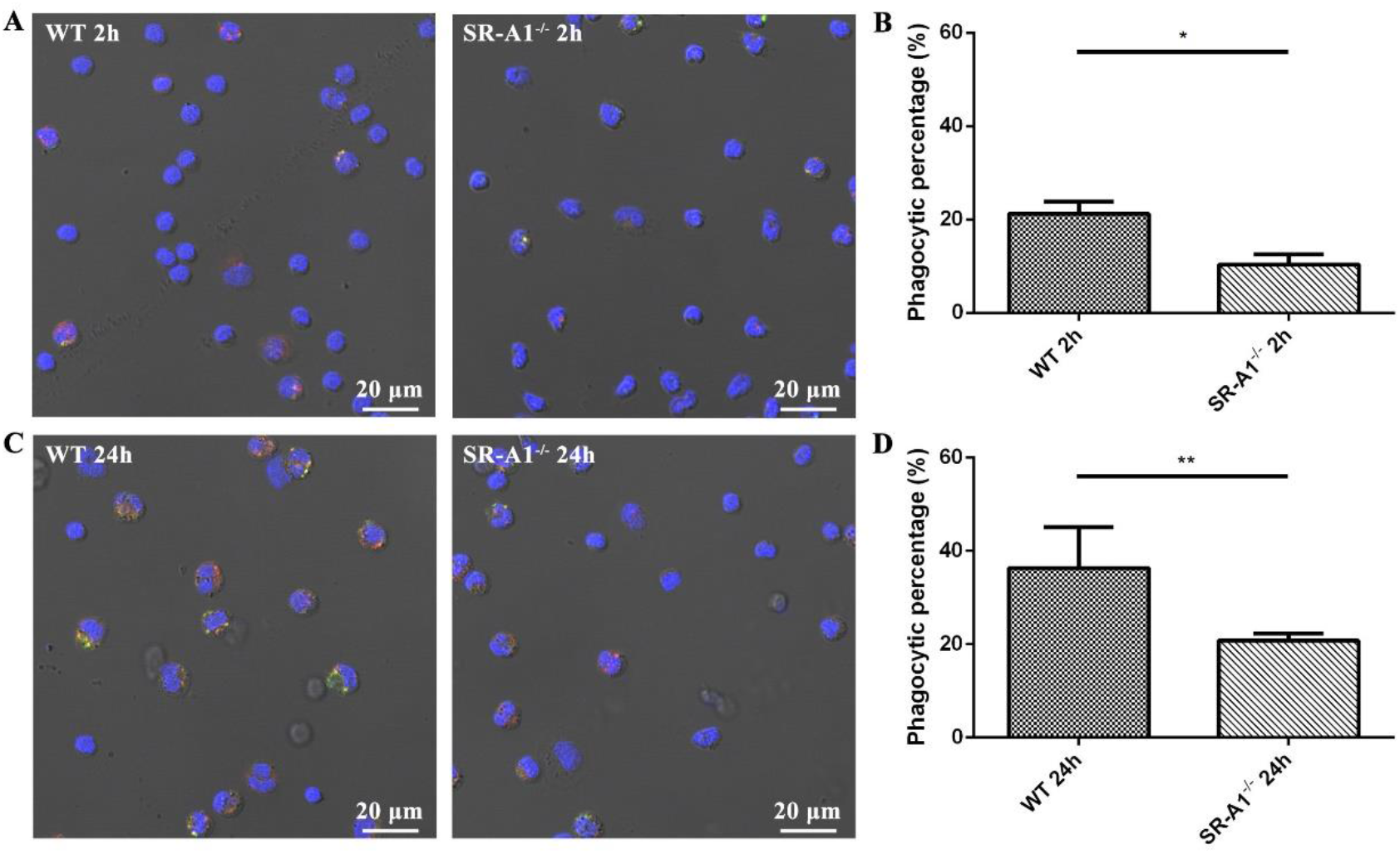
SR-A1^-/-^ PMs were deficient in phagocytosis of *L. interrogans* strain 56606v *in vivo*. WT or SR-A1^-/-^ mice were infected by ip injection of 2×10^8^ bacteria per mouse. PMs were harvested at 2 and 24 hpi, respectively. Rabbit anti-*L. interrogans* strain 56606v was used as a specific primary antibody, while FITC-conjugated or TRITC-conjugated anti-rabbit IgG as a secondary antibody were used before and after permeabilization, respectively. Confocal microscopic images showed leptospires inside (red) or outside (yellow) of PMs at 2 hpi **(A)** and 24 hpi **(C)**. Phagocytic percentages were calculated and statistically analyzed **(B, D)**. These data were expressed as the mean ± SEM from at least three experiments. *\*\*P*<*0*.*01*.

### Leptospiral LPS is the potent ligand for murine SR-A1

It was reported that LPS could bind SR-A1 and is subsequently internalized in macrophages (Hampton, Golenbock, Penman, Krieger, & Raetz, 1991), which was a useful candidate ligand for SR-A1 uptake of leptospires. However, leptospiral LPS has been shown to be distinctly different from that of other Gram-negative bacteria, both structurally and functionally (Werts et al., 2001). To test whether leptospiral LPS was responsible for SR-A1 binding and phagocytosis of leptospires, quantified microbeads coated with purified leptospiral LPS or BSA (Supplementary Fig. 5) were prepared and then cocultured with macrophages. Confocal microscopy was performed to detect the phagocytic capacity of coated beads. WT BMDMs had approximately 50% more in the uptake/bind of LPS-coated beads than SR-A1^-/-^ BMDMs (Fig. 5A, B). There were also no differences in the uptake/bind of the LPS-coated beads and BSA-coated beads in SR-A1 deficient cells. (Fig. 5A, B). Similar results were obtained that HEK293T expressing SR-A1 bound more LPS-coated beads than BSA-coated beads (Fig. 5C, D). No differences were found with BSA-coated beads bound by HEK293T SR-A1 or mock cells (Fig. 5C, D). These results indicated that leptospiral LPS, as the ligand of SR-A1, can be endocytosed/binded by SR-A1.

**Fig. 5.**
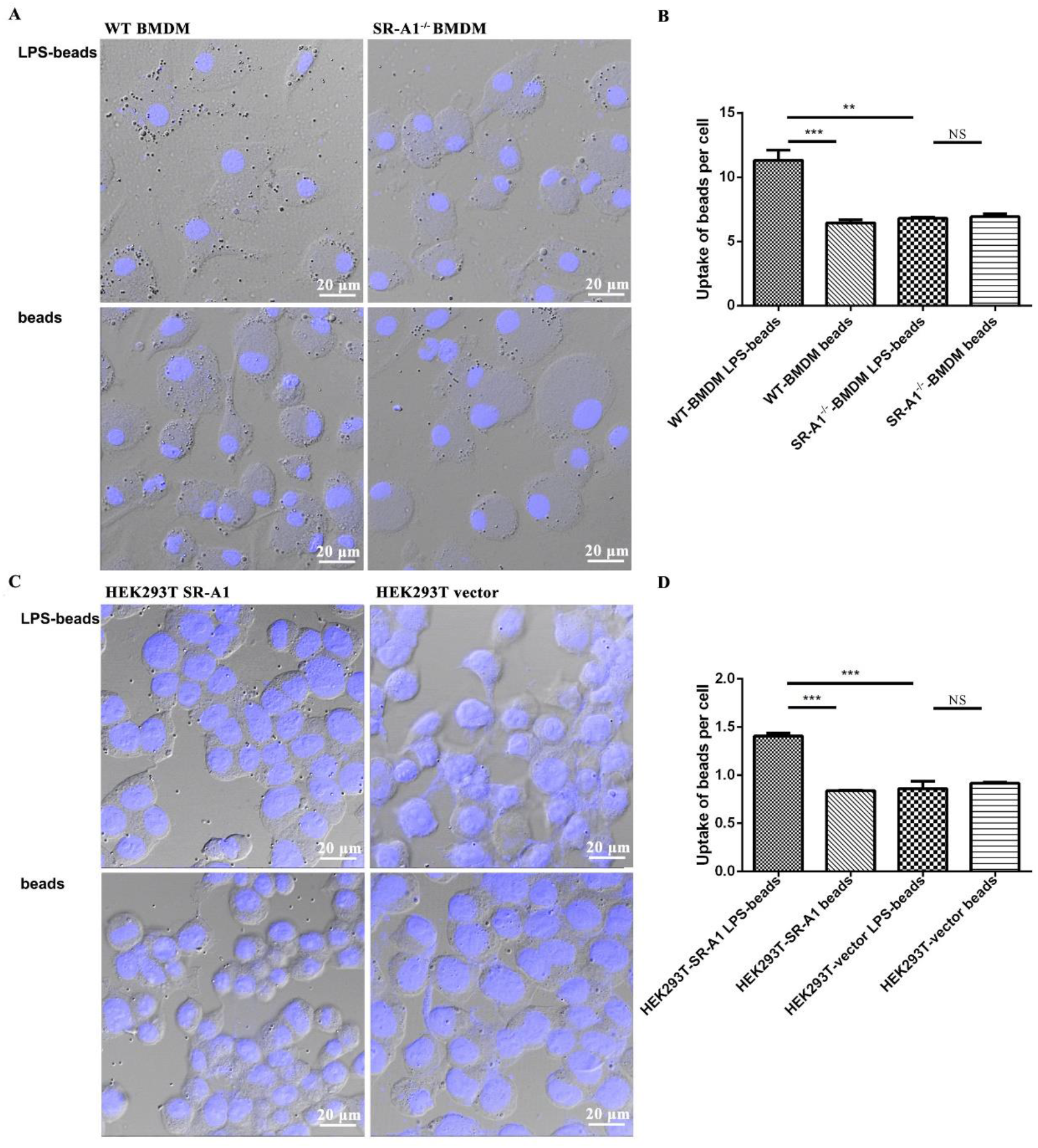
LPS of *L. interrogans* strain 56606v can be endocytosed/binded by SR-A1. BMDMs from mice **(A)** or transfected HEK293T cells **(C)** were incubated with LPS-coated or BSA-coated beads at an MOI of 10 for 1h in medium without FBS. **(B, D)** After viewing with a confocal microscope, uptake of beads per cell were calculated, respectively. These data were expressed as the mean ± SEM from at least three experiments. *\*\*P*<*0*.*01, ***P*<*0*.*001*, NS no significance.

### SR-A1^-/-^ mice were more resistant to *L. interrogans* infection compared with WT mice

It has been reported that SR-A1^-/-^ mice were more susceptible to some bacterial infections than WT mice (Arredouani et al., 2006; Ishiguro et al., 2001; Peiser et al., 2002). This is most likely due to SR-A1 mediating opsonin-independent phagocytosis of bacteria, which plays a critical role in host defense against bacterial infections. To determine whether SR-A1 also plays an important role in defense against *L. interrogans in vivo*, WT and SR-A1^-/-^ mice were infected by intraperitoneal injection of 2×10^8^ *L. interrogans* strain 56606v, and the pathological lesions were examined at 1, 3 and 5 days post-infection. None of the WT or SR-A1^-/-^ mice died from the infection during the study. Strikingly, infected WT mice developed more obvious jaundice and subcutaneous hemorrhage than SR-A1^-/-^ mice (Fig. 6A). Two liver serological markers, total bilirubin (TBIL) and aspartate transaminase (AST), were found elevated in infected mice with significantly higher in WT than in SR-A1^-/-^ mice. In contrast, the kidney serological marker creatinine (CREA) remained at normal levels in both WT and SR-A1^-/-^ mice (Fig. 6B).

**Fig. 6.**
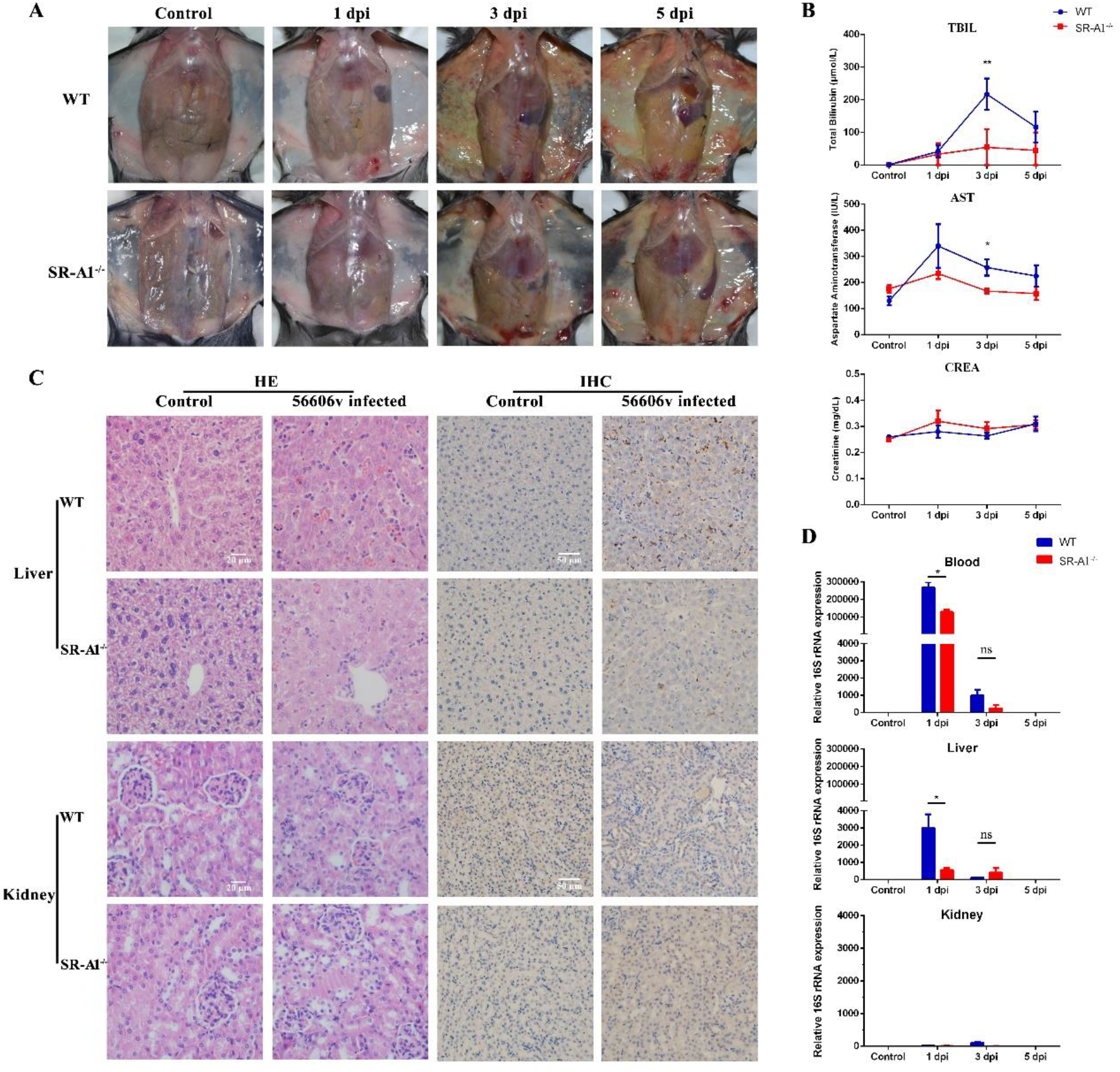
SR-A1^-/-^ mice were more resistant to *L. interrogans* infection as compared to WT mice. WT or SR-A1^-/-^ mice were infected by IP injection of 2×10^8^ bacteria per mouse. **(A)** Gross observation of jaundice and hemorrhage in abdominal and subcutaneous tissue was illustrated. **(B)** The levels of serum total bilirubin (TBIL), aspartate transaminase (AST) and creatinine (CREA) on behalf of liver and kidney function were measured by UniCel DxC 800 Synchron autoanalyzer. **(C)** HE staining in different organs was performed for histopathological analysis, and the leptospires in organs were detected by immunohistochemistry. **(D)** The leptospiral loads in the blood, liver and kidney of mice at 1-5 dpi were determined through qPCR method of bacterial 16S rRNA. The results were calculated by mean ± SEM from three mice per time point and were representative of three independent experiments. \**P*<0.05, \*\**P*<0.01, NS no significance between WT and SR-A1^-/-^ mice.

H&E staining revealed a reduced severity of acute leptospirosis in the liver of SR-A1^-/-^ mice compared with that of WT mice (Fig. 6C). The loss in the liver architecture of WT mice was observed along with hepatocyte focal necrosis, hemorrhage and Kupffer cell hyperplasia. But only a mild focal hemorrhage was observed in SR-A1^-/-^ mice. There were no severe lesions in the kidney tissues of both mice (Fig. 6C). None of the pathological changes was observed in PBS-inoculated control animals.

The leptospiral loads were determined by estimates of the copy number of *Leptospira* 16S rRNA in blood and tissues. A significantly lower bacterial burden was observed at 1 dpi in the blood and liver of the infected SR-A1^-/-^ mice than in WT mice (Fig. 6D). Leptospires were also detected in different organs by immunohistochemistry (Fig. 6C). Consistent with the 16S rRNA detection, SR-A1^-/-^ mice showed lower bacterial burdens in the liver than in those of WT mice (Fig. 6C). There was no difference in leptospiral loads in the kidney tissues between infected SR-A1^-/-^ and WT mice.

In the detection of cytokines, we found that the expression of iNOS and pro-inflammatory factors such as IL-1β and TNFα was significantly reduced in the blood and liver of SR-A1^-/-^ mice (Fig. 7). IL-6 expression *in vivo* was not affected by SR-A1. There was no severe inflammation in kidney tissues from either group (data not shown) and were consistent with pathology experiments in Fig. 6. Our results were consistent with a previous report that severe hemorrhaging caused by *Leptospira* is related to the overexpression of inflammatory cytokines and mediators (Xia et al., 2017).

**Fig. 7.**
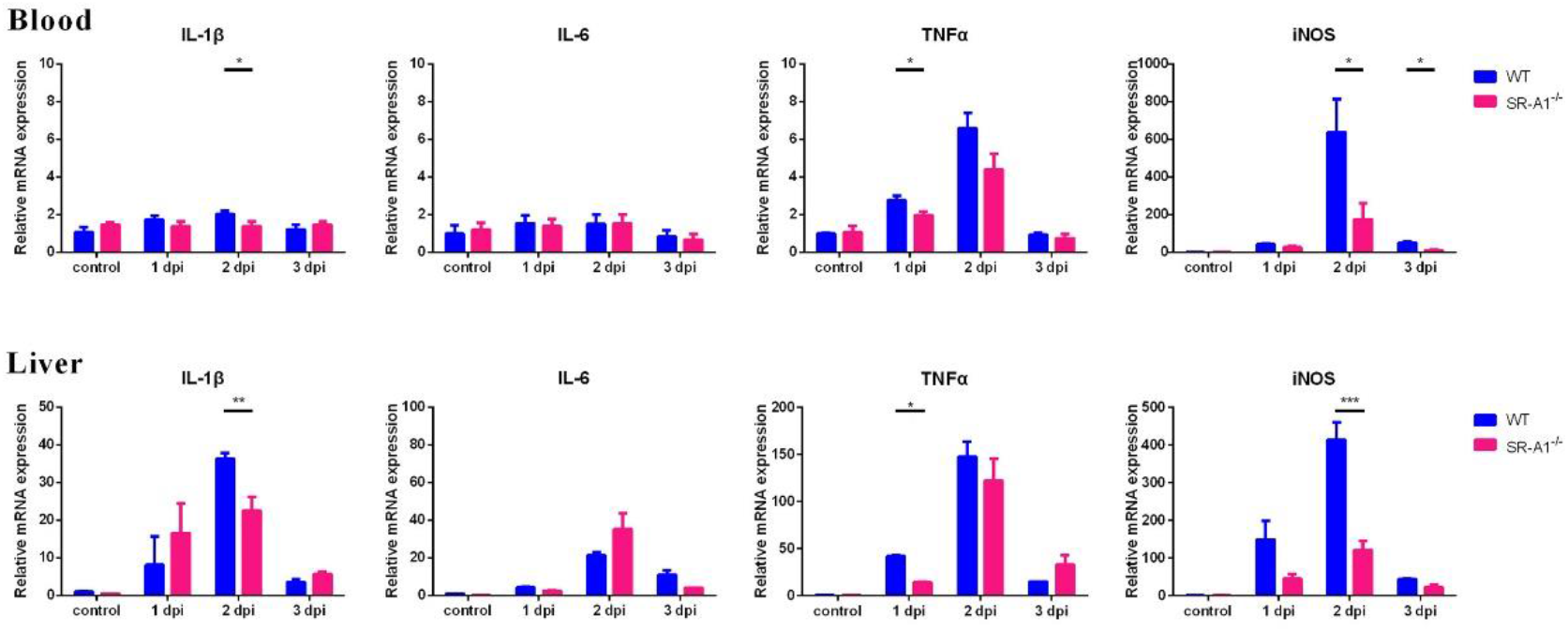
SR-A1^-/-^ mice illustrated reduced inflammatory response to *L. interrogans* infection as compared to WT mice. WT or SR-A1^-/-^ mice were infected by IP injection of 2×10^8^ *L. interrogans* strain 56606v per mouse. Cytokines expression in the blood and liver of mice at 1-3 dpi were determined through qPCR method. The results were calculated by mean ± SEM from three mice per time point, and were representative of three independent experiments. \**P*<0.05 between WT and SR-A1^-/-^ mice.

### SR-A1 increase live *L. interrogans* loads in macrophages *in vitro*

Decreased *L. interrogans* burden in SR-A1^-/-^ mice might seem counterintuitive since SR-A1 increases *Leptospira*’*s* uptake by macrophages. SR-A1^-/-^ mice cleared *Leptospira* from the liver and blood more efficiently compared with WT mice, suggesting that SR-A1 contributes to *Leptospira* infection pathophysiology. Previous studies reported that *Leptospira* could survive and replicate in macrophages and even be transported to organs by infected macrophages (Toma, Okura, Takayama, & Suzuki, 2011). In this study, immunohistochemical results showed that macrophage recruitment of *Leptospira* infected WT mice in liver tissue was significantly higher than that of SR-A1^-/-^ mice (Supplementary Fig. 7), which was consistent with the previous findings that SR-A1 might influence macrophage recruitment (Cholewa, Nikolic, & Post, 2010). Thus, we postulate that SR-A1 might enhance the live leptospires load in macrophages and contribute to leptospiral dissemination in organs.

To test this hypothesis, we performed the *in vitro* experiment to detect leptospiral survival within macrophages. We found a small proportion of ingested leptospires survived in macrophages at 72 hpi (Supplementary Fig. 6), although most internalized *Leptospira* were killed at 48 hpi. The live leptospires load was higher in WT macrophages than in SR-A1^-/-^ macrophages at 72 hpi (Supplementary Fig. 6). These results indicated that SR-A1 increases the live leptospires load in macrophages and might promote the dissemination of *Leptospira* in organs.

### SR-A1 is essential for intracellular receptor-mediated cytokine and inflammation mediator secretion in macrophages infected by *Leptospira in vitro*

I*n vivo* studies showed that WT mice had higher *Leptospira* loads and increased inflammatory responses than those of SR-A1^-/-^ mice. Exacerbated inflammatory responses in WT mice might be due to the consequence of high *Leptospira* loads or the inflammatory process triggered by SR-A1. Prior studies suggested that pathogen internalization via SR-A1 prevents sustained sensing and response by surface TLRs, increasing the intracellular immune responses through intracellular receptors, such as NOD1, NALP3 and TLR3 (Mukhopadhyay et al., 2011). To evaluate whether SR-A1 has a direct role in modulating inflammatory responses, we measured NO and cytokines expression using WT and SR-A1^-/-^ PMs infected with *Leptospira in vitro*. We performed ELISA test of supernatants from PMs stimulated with *L. interrogans* as MOI 100 at 1 dpi *in vitro*. NO and IL-1β expression levels were significantly decreased in SR-A1^-/-^ cells (Fig. 8A), consistent with cytokine expression *in vivo* (Fig. 7). However, IL-6 and TNFα expression *in vitro* were not affected by SR-A1 (Fig. 8A). The transcriptional level of TNFα, IL-6, IL-1β and iNOS mRNA in *Leptospira* infected macrophages were also detected. The results showed that SR-A1^-/-^ macrophages produced significantly less iNOS mRNA compared with WT macrophages (Fig. 8B). However, the mRNA levels of IL-1β, IL-6 and TNFα were not affected by SR-A1 (Fig. 8B).

**Fig. 8.**
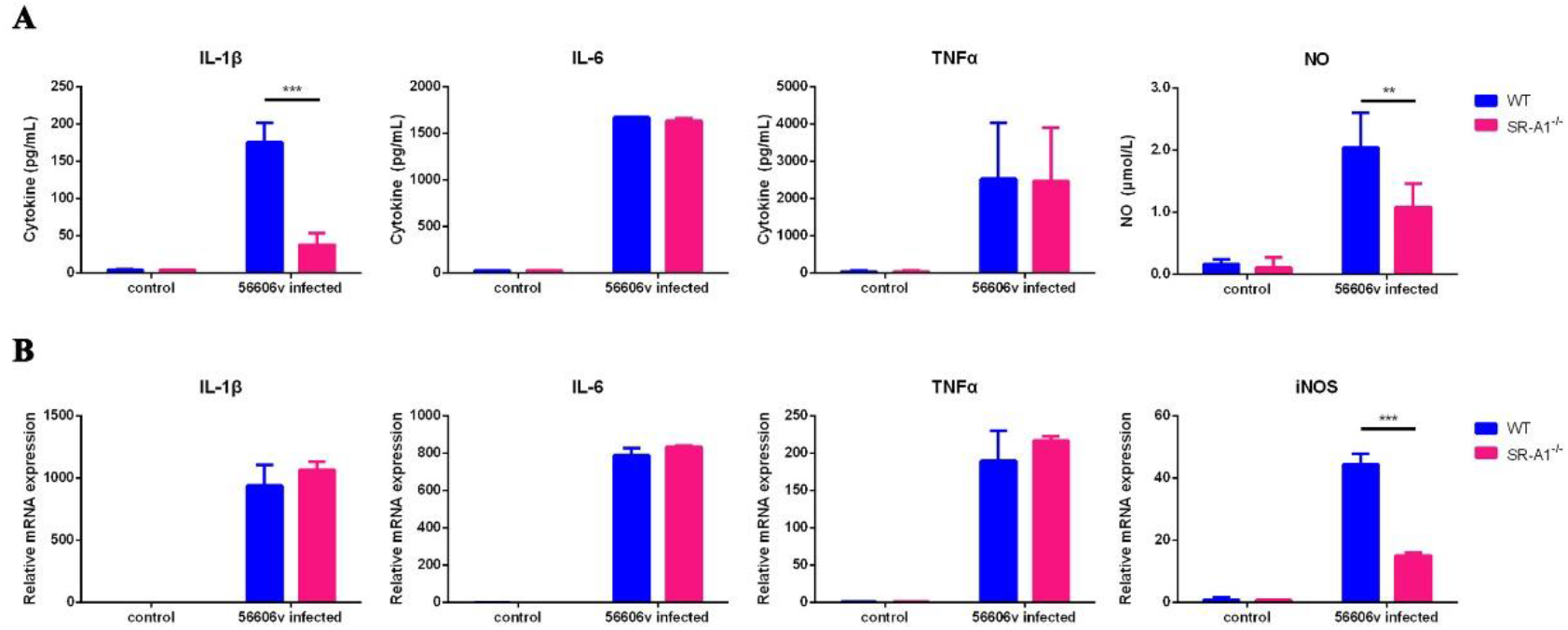
SR-A1 enhance the intracellular receptor activation in *L. interrogans* infected macrophages *in vitro*. PMs from WT or SR-A1^-/-^ mice were infected by *L. interrogans* strain 56606v as MOI=100 *in vitro*. **(A)** The cytokine level of the supernatant at 1 dpi were quantified by ELISA kits from R&D Systems, and contemporaneous NO levels was tested immediately via the Griess reaction. (B) Cytokines and iNOS expression of PMs at 3 hpi were quantified through the RTqPCR method. These data were expressed as the mean ± SEM from at least three experiments. *\*\*P*<*0*.*01, ***P*<*0*.*001*.

Pathogen ligands could activate macrophages by binding to TLR4 and triggered two major pro-inflammatory signaling pathways, including MyD88-dependent pathways triggered at the plasma membrane and TRIF-dependent pathways in endosomes (Coburn et al., 2020). The induction of TNFα and IL-6 was mainly MyD88-dependent, and the results indicated that surface receptor-mediated cytokines induced by *Leptospira* are unaffected by SR-A1. On the other hand, the production of NO, which was mostly dependent on the intracellular receptors, such as the TRIF adaptor or TLR3, was upregulated by SR-A1. In addition, the ELISA results also showed SR-A1 upregulated IL-1β secretion. IL-1β secretion requires TLR-mediated induction of pro-IL-1β, followed by activation of the NALP3 inflammasome and cleavage of pro-IL-1β to mature IL-1β (Godoy, Murgas, Tichauer, & Von Bernhardi, 2012). IL-1β mRNA levels, representative of TLR-mediated induction, were not affected by the presence of SR-A1, indicating the increased IL-1β cytokine secretion was mainly due to the activation of NALP3-mediated cleavage of pro-IL-1β to mature IL-1β. Our results suggested that *Leptospira* internalization via SR-A1 is essential for intracellular receptor-mediated inflammatory cytokine and mediator secretion.

## Discussion

Scavenger receptor A1, known as the macrophage scavenger receptor that binds and traffics a variety of microbial ligands, plays multiple roles in inflammation, innate immunity and host defense (Areschoug & Gordon, 2009). SR-A1 is a phagocytic receptor for a variety of gram-positive and gram-negative bacteria (Dunne et al., 1994; Peiser et al., 2002; Peiser et al., 2000). However, some bacteria, such as *Escherichia coli* K12, *Pseudomonas aeruginosa* PA14, *Streptococcus agalactiae* and *Streptococcus pyogenes*, were described as being uptake by macrophages through an SR-A1-independent way (Amiel, Acker, Collins, & Berwin, 2009; Domingo-Gonzalez et al., 2013). The surface M protein and the sialylated polysaccharide capsule of streptococci prevent the SR-A1 mediated phagocytosis by macrophages (Areschoug, Waldemarsson, & Gordon, 2008). Unlike other typical gram-negative bacteria, *Leptospira* is a bacterium with unique evolutionary and structural characteristics. Our understanding of the receptors involved in the process of *Leptospira’s* phagocytosis is still minimal. Several host receptors such as CR3 receptor and β2 integrin have been proved to be involved in the phagocytosis of *L. interrogans* (Cinco et al., 2002; Cosate et al., 2016), while some PRRs, like the mannose receptors (MRs) as shown above in our study, are less likely to participate in this process.

This study demonstrates that SR-A1 is a primary nonopsonic phagocytic receptor for *Leptospira* on murine macrophages. We utilized SR-A1 inhibitor, SR-A1 blocking antibody, SR-A1^-/-^ murine macrophages, and SR-A1 overexpressed cells to examine the interaction of SR-A1 and *Leptospira*. The application of SR-A1 inhibitor poly I, lacks specificity in binding SR-A1, while the SR-A1^-/-^ macrophages induce compensatory up-regulation of alternative receptors, causing misinterpretation of the results. To avoid these issues, we used the SR-A1 specific antibody to block the SR-A1 binding function. Our results showed that *Leptospira* phagocytosis ratio by SR-A1^-/-^ macrophages are down-regulated to appropriately 50%. A similar result was also obtained when WT macrophages were treated with SR-A1 blocking antibody. These results indicate that SR-A1 plays a vital role in phagocytosis *in vitro*.

SR-A1^-/-^ mice were used to investigate SR-A1 phagocytosis function *in vivo*. After intraperitoneal infection, invading *Leptospira* can be opsonized by natural antibodies and complement in the plasma, promoting their phagocytosis through CR3 receptors. Our results showed that WT PMs have higher phagocytic rates than SR-A1^-/-^ PMs (Fig. 4) and the phagocytic function of SR-A1 was not concealed by the effect of opsonic receptors in the peritoneal fluid of experimental infected animals, suggesting that SR-A1 also plays a significant role in the phagocytosis of *Leptospira in vivo*.

We extended our studies to focus further on bacterial ligands for SR-A1. Previous studies showed that SR-A1 could interact with various bacterial components, including LPS, lipoteichoic acid, and polyribonucleic acids (Dunne et al., 1994; Hampton et al., 1991; Zhu, Reich, & Pisetsky, 2001). *Leptospira* ligand direct binding to SR-A1 is presently unknown, and the best candidate ligand for SR-A1 recognition is LPS. Several reports have shown that SR-A1 can bind LPS via recognition of the lipid A molecule (Ashkenas et al., 1993; Hampton et al., 1991). However, LPS is not always the ligand of SR-A1 during its interaction with different bacteria. Peiser and his colleagues showed that SR-A1 recognized *Neisseria meningitidis* independently of LPS (Peiser et al., 2006). *Escherichia coli* K12, with a rough-form LPS, was described as being uptake by macrophage via an SR-A1 independent mechanism (Peiser et al., 2000). Leptospiral LPS has been revealed to have distinct structural and functional features, which are different from those of enterobacterial species (Que-Gewirth et al., 2004; Werts et al., 2001). Therefore, the hypothesis of leptospiral LPS as a ligand for SR-A1 is worth further study. We utilized beads coated with leptospiral LPS to assess macrophage phagocytic capacity with or without SR-A1 receptors. The results showed that the enhanced phagocytic capacity for SR-A1 expressed cells relied on the presence of LPS (Fig. 5). The results suggested leptospiral LPS is the ligand of SR-A1.

Earlier works have suggested that the contribution of SR-A1 to host defense of bacterial infection varied with the specific strain (Ishiguro et al., 2001; Sever-Chroneos, Tvinnereim, Hunter, & Chroneos, 2011). It was shown previously that SR-A1^-/-^ mice are more susceptible to *Listeria monocytogenes, Neisseria meningitidis* and *Streptococcus pneumonia* infection than WT (Arredouani et al., 2006; Ishiguro et al., 2001; Peiser et al., 2002). In contrast, other studies showed that SR-A1^-/-^ mice have a decreased susceptibility to infection with *Mycobacterium tuberculosis, Pneumocystis carinii* and polymicrobial sepsis (Drummond et al., 2013; Hollifield, Bou Ghanem, de Villiers, & Garvy, 2007; Sever-Chroneos et al., 2011). Although studies have clearly demonstrated that SR-A1 is important in *Leptospira* uptaken by macrophages in the peritoneal cavity, it remains to be tested *in vivo* if the high phagocytosis plays a vital role in host defense. Our acute leptospirosis murine model provides a powerful tool to clarify the question (Xia et al., 2017). Using this model, mice intraperitoneally injected with leptospires displayed high bacterial loads in blood and tissues, and severe clinical symptoms of leptospirosis. Macrophages have been indicated to be the primary infiltrating and anti-*Leptospira* phagocytes during *Leptospira* infection (Chen et al., 2017). Enhancing the phagocytic activity of macrophages seems to increase the ability to fight infection. Surprisingly, leptospiral infected WT mice presented more severe jaundice, subcutaneous hemorrhage, and higher bacteria burden in blood and tissues than that of SR-A1^-/-^ mice, suggesting that *Leptospira* can exploit SR-A1 to promote their dissemination and cause severe symptoms in the host.

Multiple pathogens have evolved strategies to regulate macrophages activation and responses. *Mycobacterium marinum* and *Mycobacterium leprae* were reported taking advantage of macrophages as vehicles for bacterial dissemination (Clay et al., 2007; Madigan et al., 2017). Clay *et al*. showed that while depleting macrophages led to higher *Mycobacterium marinum* burdens and increased host death, it also decreased the dissemination of pathogens into deeper tissue (Clay et al., 2007). This was referred to as a “dichotomous role” of macrophages. In case of *L. interrogans*, previous studies using the zebrafish embryos model confirmed that infected macrophages participate in *L. interrogans* dissemination (Davis, Haake, & Ramakrishnan, 2009). Although *L. interrogans* is usually considered to be extracellular pathogens, they can survive and even replicate in murine macrophages (Li et al., 2010; Toma et al., 2011). Macrophages were capable of killing opsonized leptospires effectively but had little bactericidal activity against nonopsonized *L. interrogans*, and intact leptospires could be found in the cytosol (Banfi, Cinco, Bellini, & Soranzo, 1982; Cinco, Banfi, & Soranzo, 1981). In this study, the results of the survival of *L. interrogans* in peritoneal macrophage showed that a small proportion of ingested *L. interrogans* survived in macrophages 72 hours post-infection, and live *L. interrogans* loads were higher in WT macrophages than in SR-A1^-/-^ macrophages (Supplementary Fig. 6). This may be due to the contribution of SR-A1-related nonopsonic phagocytosis. Besides, SR-A1 was reported to have an inhibitory effect on autophagy in macrophages (Huang et al., 2014). Higher *L. interrogans* load in WT macrophage might also be related to the inhibition of autophagy by SR-A1, with the consequences of intracellular bactericidal activity diminished. Our findings suggest that infected macrophages might participate in leptospiral dissemination, as a higher *Leptospira* load in macrophage following a higher *Leptospira* burden in organs. Another characteristic of SR-A1, cell adhesion and surface localization, which are essential for macrophage recruitment to the sites of tissue infected (Cholewa et al., 2010), might be partly responsible for the higher bacteria load in the tissue of WT mice. In this study, WT mice had more macrophage recruitment in liver tissue than that of SR-A1^-/-^ mice (Supplementary Fig. 7), and further suggested that infected macrophages might exploit SR-A1 to promote leptospire spread to target organs. However, our suspected mechanism was different from previous studies reporting that phagocytes depleted by clodronate liposomes led to enhanced leptospiral burdens in the acute phase (Ferrer et al., 2018). One likely explanation for this difference might be that infected macrophages are not the sole mechanism for dissemination. Elucidation of the comprehensive mechanisms for *Leptospira* dissemination will need further study.

In this study, WT mice presented more severe symptoms and tissue damage than SR-A1^-/-^ mice. Our previous study suggested that severe hemorrhage is more related to the overexpression of iNOS and proinflammatory cytokines than at the higher burden of *Leptospira* (Xia et al., 2017). To analyze the possible correlation between the SR-A1 and inflammatory response, we evaluated the TNFα, IL-6, IL-1β and iNOS level in WT and SR-A1^-/-^ mice. We observed markedly higher levels of TNFα, IL-1β and iNOS mRNA in the liver and blood of WT mice compared to SR-A1^-/-^ mice in early stages and correlation with severe symptoms (Fig. 7). *In vitro* study using peritoneal macrophages stimulated with *Leptospira* showed similar results with higher cytokines produced by WT cell compare with that of SR-A1^-/-^ macrophages (Fig. 8). It seems that an increased inflammatory response was not only because of the higher *Leptospira* burden in the host but also due to SR-A1 activity.

In addition to its scavenging function, SR-A1 modulates inflammatory responses. However, the mechanism of how SR-A1 regulates inflammation responses is still unclear. Some contradictory results were found by several studies suggesting both pro-inflammatory and anti-inflammatory roles of SR-A1 (Godoy et al., 2012; Haworth et al., 1997; Hollifield et al., 2007; Mukhopadhyay et al., 2011; Yu et al., 2011). One theory proposed by Subhankar *et al*. is that SR-A1 attenuates TLR4-driven proinflammatory cytokine responses by endocytic scavenging of LPS from the extracellular environment, whereas SR-A1 enhances intracellular immune responses through intracellular receptors, such as NOD1, NALP3 and TLR3 (Mukhopadhyay et al., 2011). In the case of *L. interrogans* strain 56606v, our previous work showed this strain to induce low levels of LPS mediated endotoxic activity (Xia et al., 2017), which suggested negligible potential effects on MyD88-dependent pro-inflammatory signaling pathways due to clearance of LPS residues from the cell surface via SR-A1 receptor. On the other hand, SR-A1-mediated *Leptospira* internalization and intracellular survival tend to cause robust intracellular receptors’ responses. This theory is consistent with our results in Figure 8, showing that the secretion of IL-1β and NO, mediated by NALP3 and TRIF in endosomes, respectively, were increased in WT PMs compared to SR-A1^-/-^ PMs. We conclude that *Leptospira* internalization via SR-A1 is essential for intracellular receptor-mediated inflammatory cytokine and mediator secretion and further causes tissue damage.

Pathogens have evolved numerous complex mechanisms to exploit loopholes in the immune system to gain advantages over host immune defenses. Understanding the processes that pathogens manipulate immune responses are critical for controlling the infection. Our study reveals that although the host is beneficial in the uptake of *Leptospira* by macrophages, SR-A1 might be exploited by *Leptospira* to promote bacterial dissemination and inflammatory responses, which causes more severe infections in the host. Leptospiral strategies to take advantage of SR-A1 to establish severe infection are depicted in Fig. 9. This study suggests the potential of utilizing SR-A1 inhibiter in the treatment of leptospires infection.

**Fig. 9.**
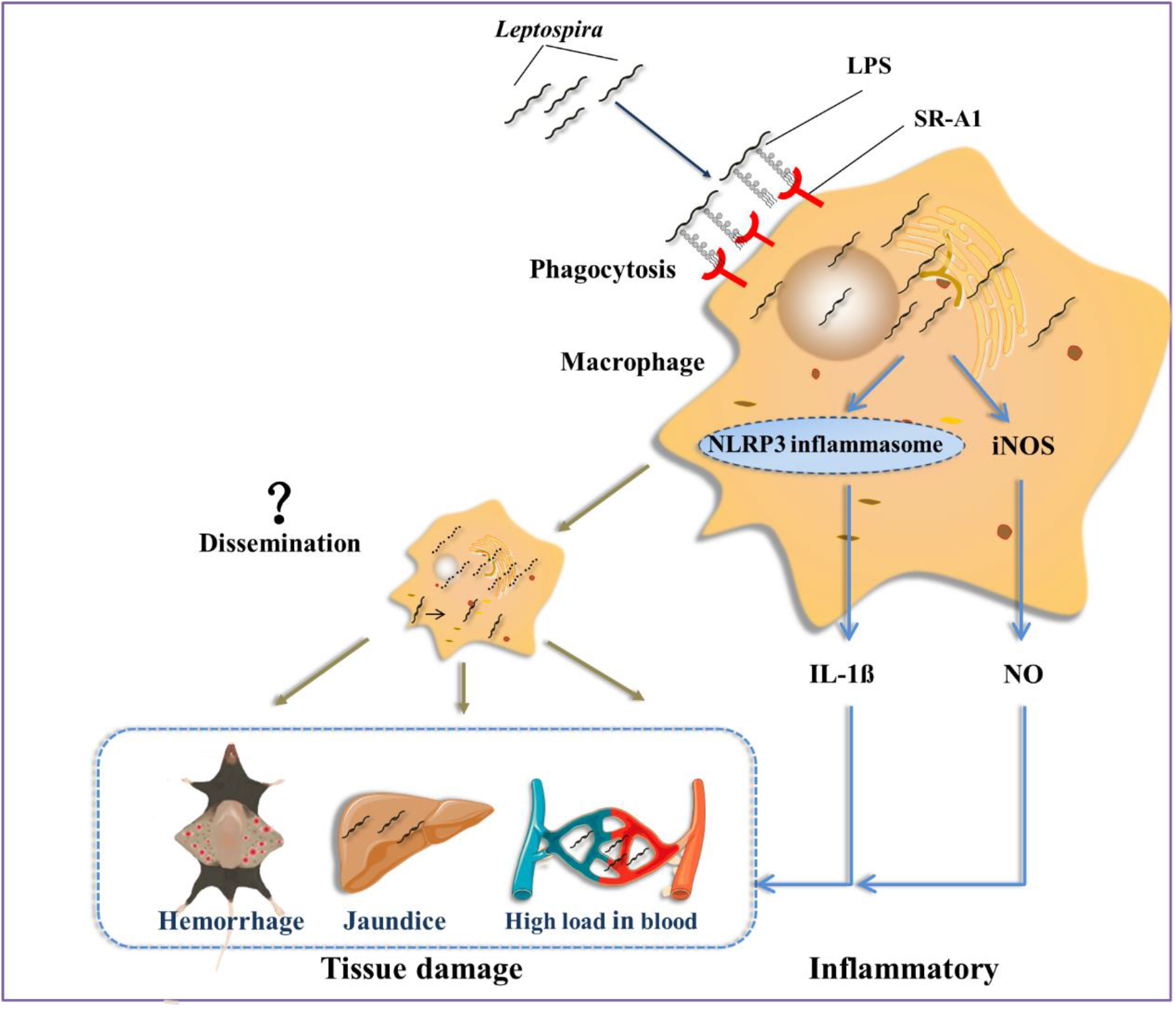
Strategies employed by *Leptospira* to take advantage of SR-A1 to establish severe infection. In *Leptospira*-infected mice, macrophages play a controversial role in controlling leptospires in the initial stage of infection. SR-A1 on macrophages binds LPS of *Leptospira*, mediating the phagocytosis of *Leptospira*. The majority of internalized *Leptospira* were killed, whereas a small proportion of ingested *Leptospira* survived in macrophages. SR-A1 mediated phagocytosis increased *Leptospira* load in macrophages and consequently might enhance the leptospiral dissemination in organs. In parallel, high *Leptospira* loads in macrophages may trigger the intracellular receptor excessive activation and enhance secretion of IL-1β and NO, which induces hemorrhage and jaundice in mice.

## Acknowledgments

We sincerely thank Dr. Chen Qi (Nanjing Medical University) for kindly providing the SR-A1^-/-^ mice. This study was supported by grants from the National Natural Science Foundation of China (grant number 81971896, 81800190, and 81471908), the Natural Science Foundation of Shanghai Program (Grant Number 19ZR1428600), National innovative research team of high-level local universities in Shanghai.

## Competing interests

The authors have declared that no competing interest exists.

## Supplemental Materials Materials and Methods

## Supplemental Materials Materials and Methods

### FITC labeling of bacterial strains

Phosphate buffered saline (PBS) washed bacteria were diluted in 1 mL of carbonate buffer solution (CBS) containing 10 μM FITC (Invitrogen, USA). The mixture for labeling was incubated for 30 min at room temperature protected from light. When using inactivated bacteria, the bacterial suspension was then fixed for 1 h at 4°C with 4% paraformaldehyde. *L. interrogans* were washed and counted in a Petroff-Hauser chamber under dark-field microscopy, and then were suspended in RPMI 1640 at a particular concentration for the experiment.

### Adhesion and phagocytosis of leptospires by flow cytometry (FCM) analysis

Fixed leptospires were pre-stained by FITC as mentioned above. Each flow tube of 1×10 ^6^ macrophages was incubated with bacteria as MOI=10 and rotated at 10 rpm in cell culture condition of 37°C. After infection, trypan blue was used to quench the fluorescence of *L. interrogans* outside the cell membrane. FCM of bacteria *in vivo* and cytoD treated cells were used to confirm quenching. Cells were harvested and fixed with 1% paraformaldehyde for FCM (Caliber, Becton Dickinson, USA) detection. A total of 10,000 cells were counted and the percentages of adhesion and phagocytosis were calculated as the number of positive cells with respect to the total events counted.

### Western blot analysis of SR-A1 overexpression

Total protein extracted from transfected HEK293T and RAW264.7 cells was resolved by SDS-PAGE and electrotransferred to polyvinylidene fluoride (PVDF) membranes (Millipore, USA), and was blocked with 2% nonfat milk (Sangon Biotech, China). SR-A1 primary antibody (rabbit monoclonal antibody, Abcam, USA) and HRP-conjugated secondary antibody (anti-rabbit IgG, Beyotime, China) were used to detect the SR-A1 expression. The blots were visualized with a chemiluminescent substrate (Thermo Scientific, USA) and exposed to ImageQuant LAS4000 (GE Healthcare, USA).

### FCM detection of cell surface overexpression of SR-A1

Cell surface SR-A1 expression in transfected HEK293T and RAW264.7 cells were detected through FCM method. Cells were harvested and fixed with 1% paraformaldehyde without permeabilization. SR-A1 primary antibody (rabbit monoclonal antibody, Abcam, USA) and Alexa Flour 647-conjugated secondary antibody (anti-rabbit IgG, Abcam, USA) were used to detect the SR-A1 expression on cell surface.

### FCM detection of leptospiral LPS coated beads

Extracted leptospiral LPS was quantified using the Limulus test. Microbeads of 1 μm diameter were coated by leptospiral LPS at 37°C for 1 h. For FCM analysis, rabbit anti-*L. interrogans* strain 56606v was used as the specific primary antibody, and FITC-conjugated anti-rabbit IgG as a secondary antibody was incubated for 30 min, respectively. After additional centrifugation for removing unbound antibody, particles were detected through FCM. A total of 10,000 microns were counted and the percentages of FITC labeled particles were calculated as the coating effect of LPS coated beads.

### Detection of survival of *L. interrogans* strain 56606v in leptospires infected WT and SR-A1^-/-^ peritoneal macrophages (PMs)

WT and SR-A1^-/-^ PMs were seeded by 1×10^6^ cells per well in a 12-well plate, and then were infected by *L. interrogans* strain 56606v as MOI=100 for 2 h *in vitro*. 100 μg/mL gentamicin was used for 1 h to kill the extracellular leptospires. Leptospiral RNA from cells of 1 hpi to 72 hpi was extracted and quantified by real-time PCR as mentioned in the main text.

### Macrophages recruitment in liver after *L. interrogans* stimulation *in vivo*

WT and SR-A1^-/-^mice were infected with 2×10^8^ leptospires via intraperitoneal route. Livers from 1 to 2 dpi of infected mice were fixed in neutral buffered 4% formaldehyde, followed by embedment of the tissues in paraffin. Sections were immunohistochemically stained for macrophage-specific F4/80 rabbit antibody of 1:600 dilution (Servicebio, China). Macrophages were counted in at least 10 HP field, and the average number of F4/80 expressing cells per HP field was calculated.

## Supplementary Table

**Supplementary Table 1.**
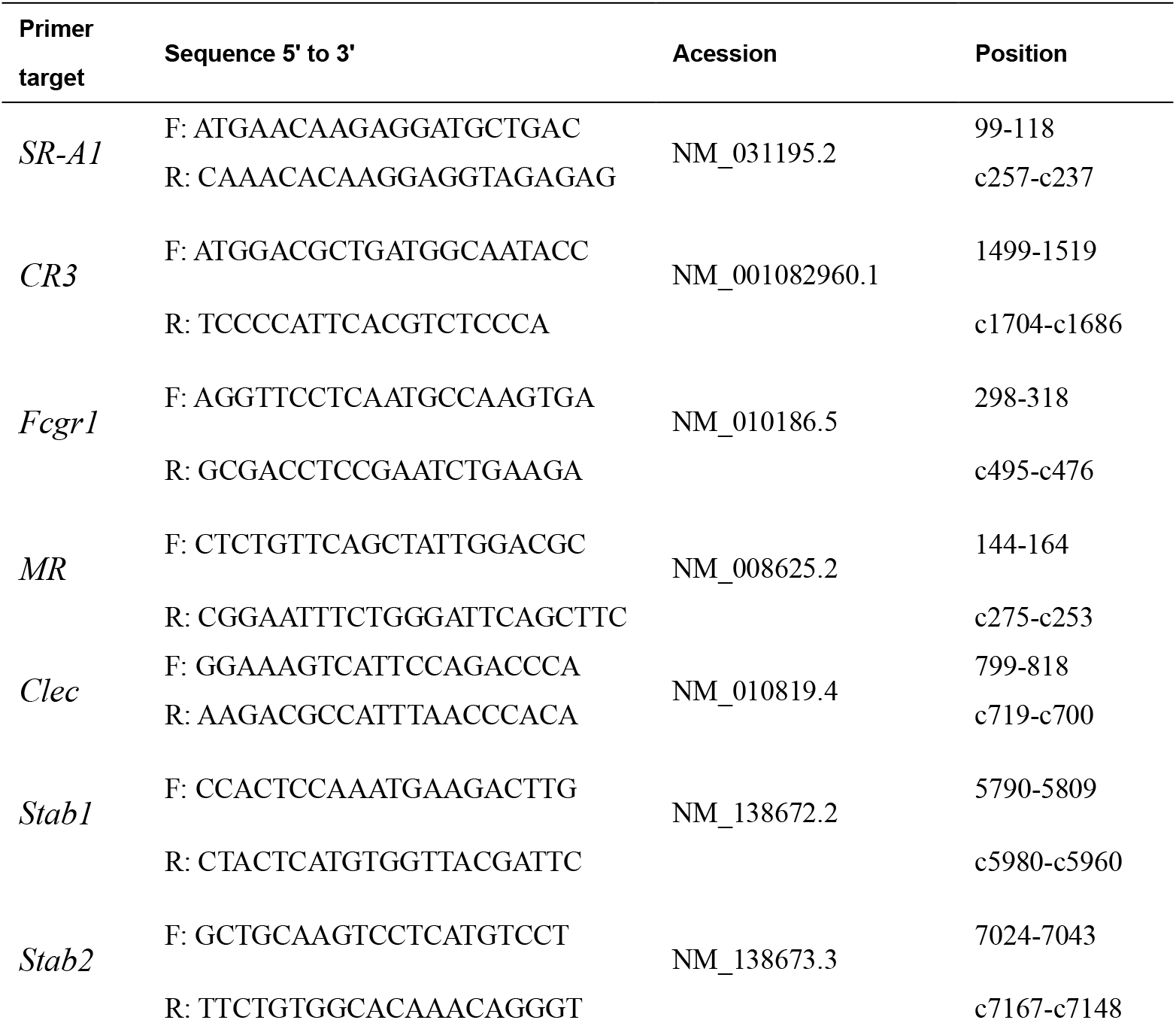
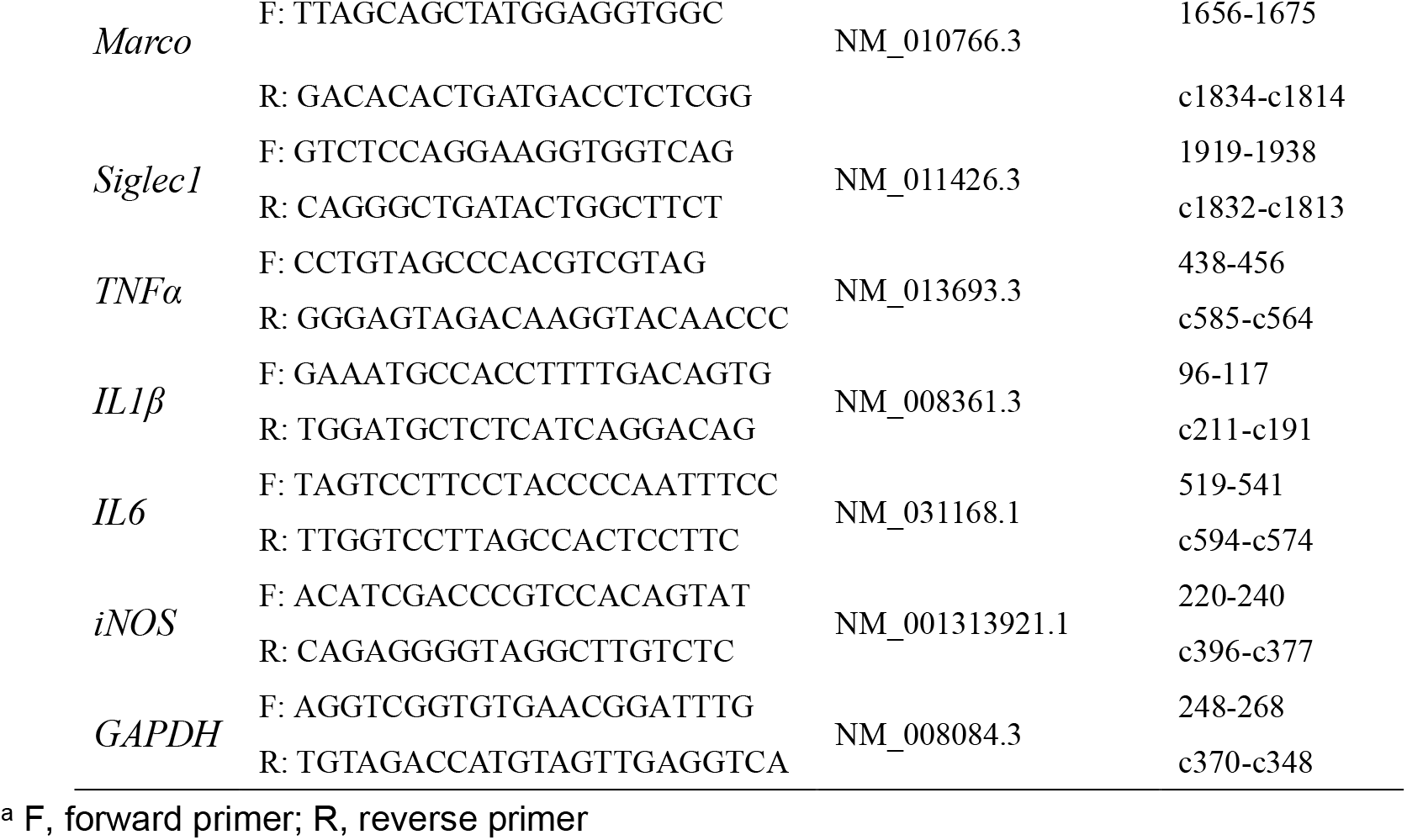
Primers used to detect receptors and cytokines expression.

## Supplementary Figures

**Supplementary Fig. 1.**
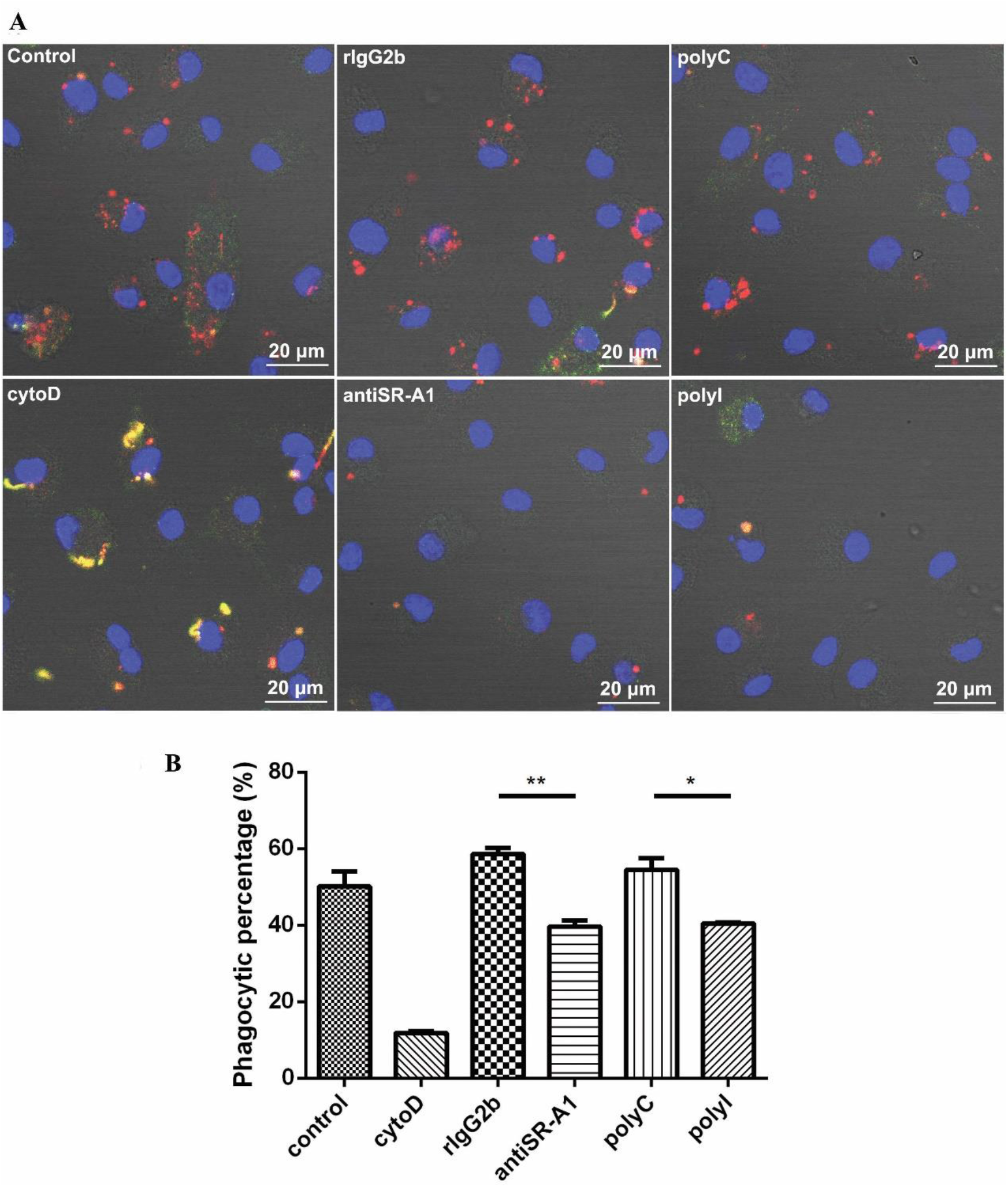
PolyI and SR-A1 monoclonal antibody (anti-SR-A1) exhibited the inhibition to phagocytosis of live *L. interrogans* strain 56606v by mouse bone marrow-derived macrophages (BMDMs). BMDMs were incubated with active *L. interrogans* strain 56606v in FBS-free medium absence or presence of cytoD (20μM), polyI (100μg/mL) and anti-SR-A1 (30μg/mL). Corresponding concentrations of polyC and rat IgG2b (rIgG2b) isotypes were added as controls. Rabbit anti-*L. interrogans* strain 56606v was treated as specific primary antibody, while FITC-conjugated or TRITC-conjugated anti-rabbit IgG as secondary antibody were used before and after permeabilization, respectively. After observations by confocal microscopy **(A)**, positive rates of BMDMs phagocytizing *Leptospira* were calculated and statistical analyzed by analysis of variance **(B)**. These data were expressed as the mean ± SEM from at least three experiments. *\*P*<*0*.*05, **P*<*0*.*01*.

**Supplementary Fig. 2.**
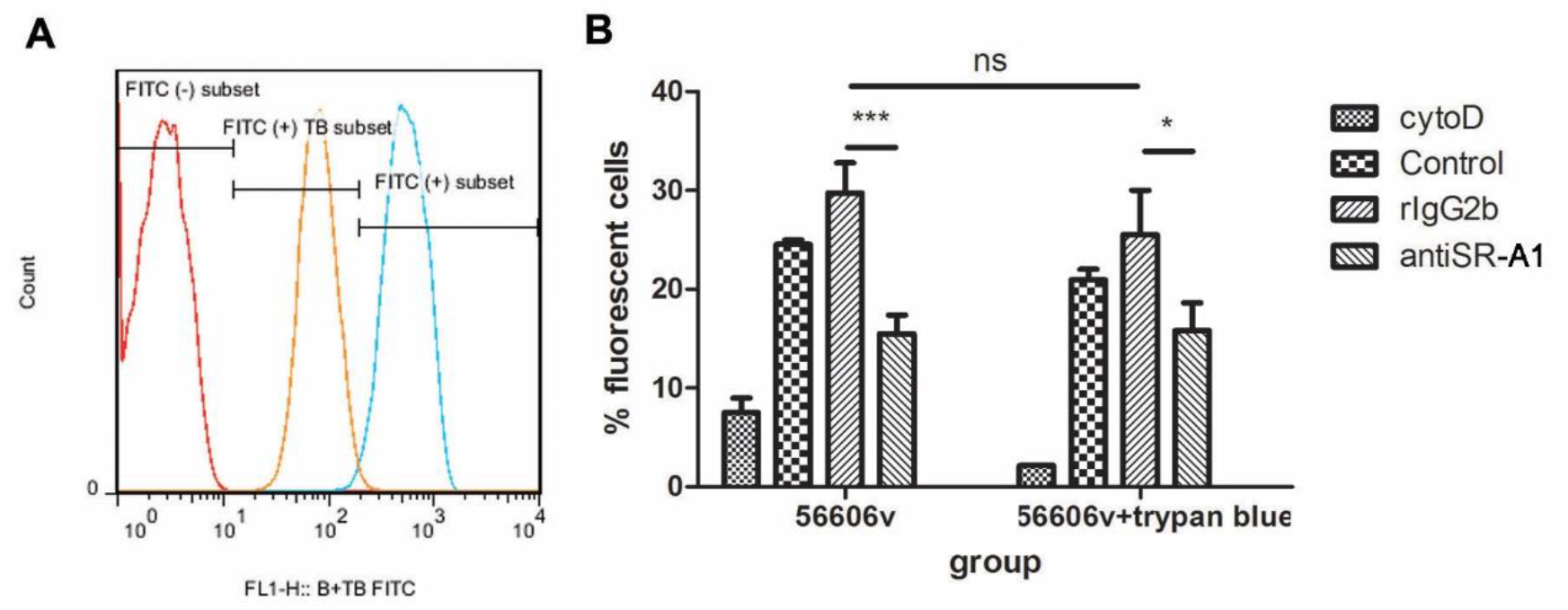
FCM exhibited the inhibition of SR-A1 inhibitor in mouse BMDMs to the phagocytosis of *L. interrogans* strain 56606v. **(A)** Inatcive leptospires were stained by FITC and quenched by trypan blue. **(B)** Macrophages pre-incubated with 20μM cytoD, 30μg/mL rIgG2b or anti-SR-A1, were co-incubated with the bacteria as MOI 10 for 30 min. Trypan blue was also used to quench the extracellular fluorescence of *Leptospira*. Percentages of phagocytosis with/without adhesion cells (trypan blue not quenched or quenched) were calculated as the number of positive cells with respect to the total events counted. These data were expressed as the mean ± SEM from at least three experiments. *\*P*<*0*.*05, ***P*<*0*.*001, ns* no significance.

**Supplementary Fig. 3.**
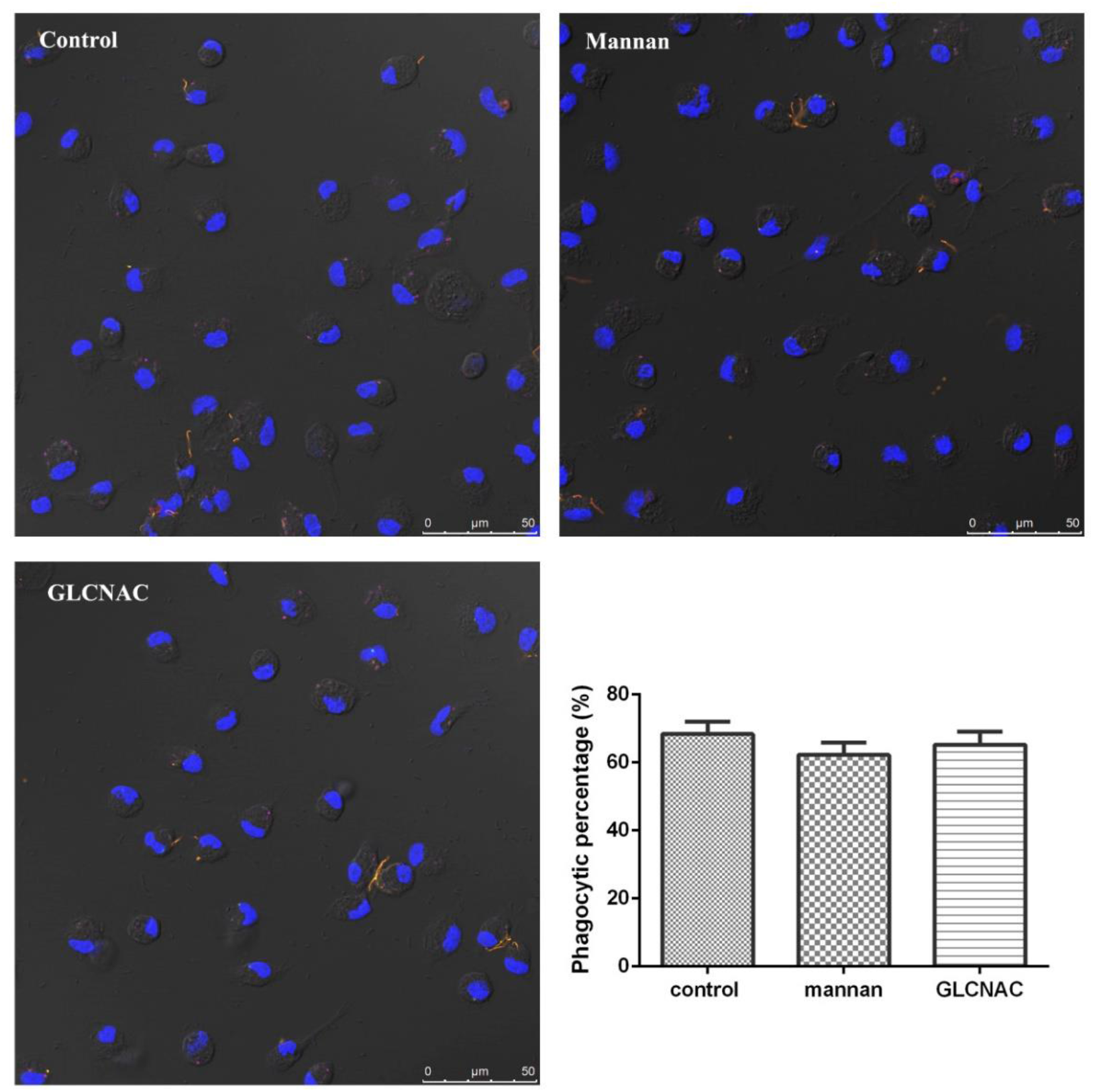
Mannan receptor inhibitor exhibited no inhibition to the phagocytosis of L. interrogans strain 56606v by mouse PMs. PMs were incubated with inactive *L. interrogans* strain 56606v in FBS-free medium absence or presence of mannan which was the competitive substance of mannan receptor. Corresponding isotype substance N-acetyl-glucosamine (GLCNAC) was added as controls. Rabbit anti-*L. interrogans* strain 56606v was treated as a specific primary antibody, while FITC-conjugated or TRITC-conjugated anti-rabbit IgG as a secondary antibody were used before and after permeabilization, respectively. After observations by confocal microscopy, positive rates of PMs phagocytizing *Leptospira* were calculated and statistically analyzed. These data were expressed as the mean ± SEM from at least three experiments.

**Supplementary Fig. 4.**
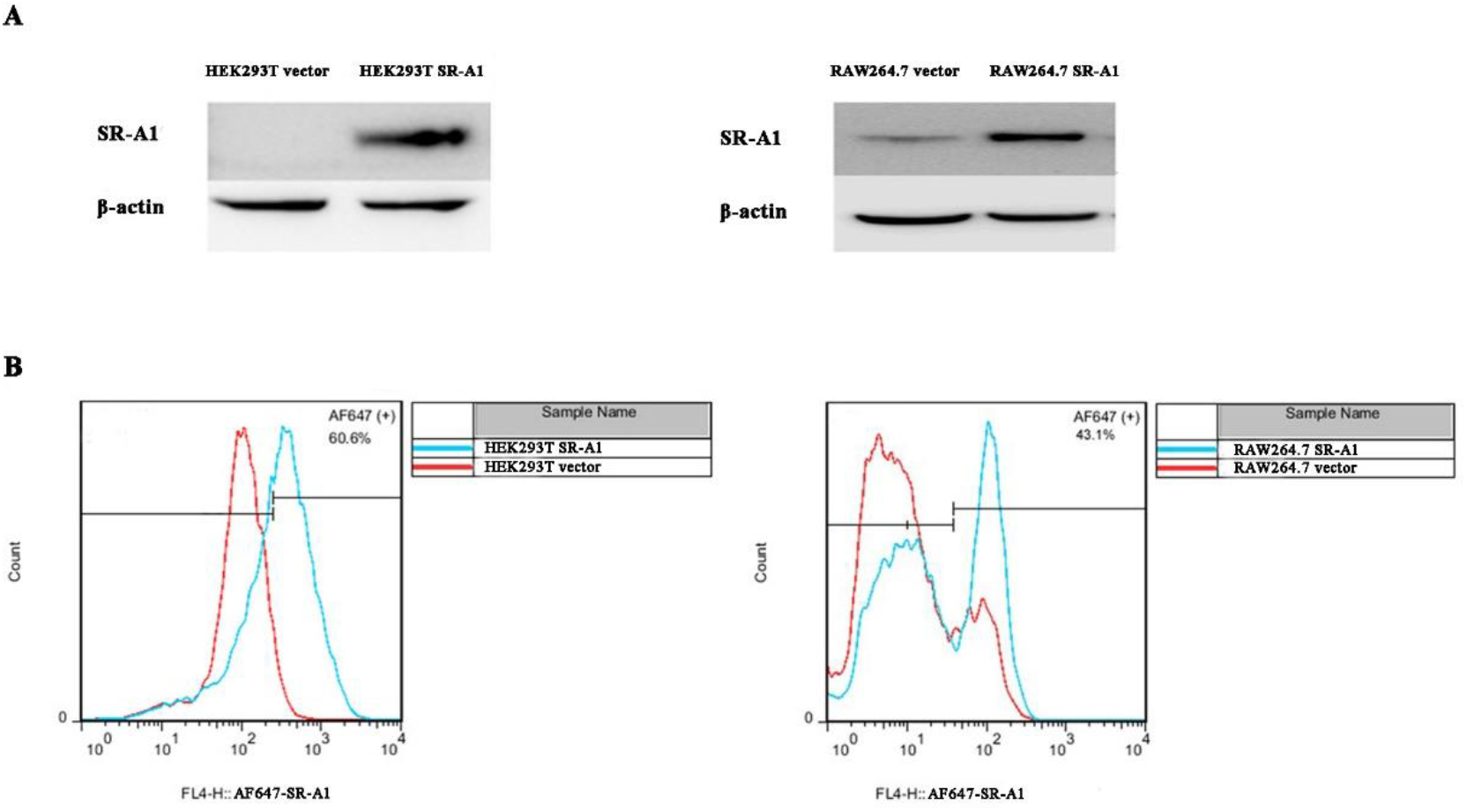
Verification of SR-A1 overexpression. **(A)** SR-A1 overexpression in transfected HEK293T and RAW264.7 cells were detected through Western blot method by SR-A1 primary antibody and HRP-conjugated secondary antibody. Relative protein expression was visualized on the blots. **(B)** Cell surface SR-A1 expression in transfected HEK293T and RAW264.7cells were detected through FCM method by SR-A1 primary antibody and Alexa Flour 647 (AF647) - conjugated secondary antibody. Positive rates of overexpressed cells verifying successful SR-A1 overexpression on cell surface were displayed.

**Supplementary Fig. 5.**
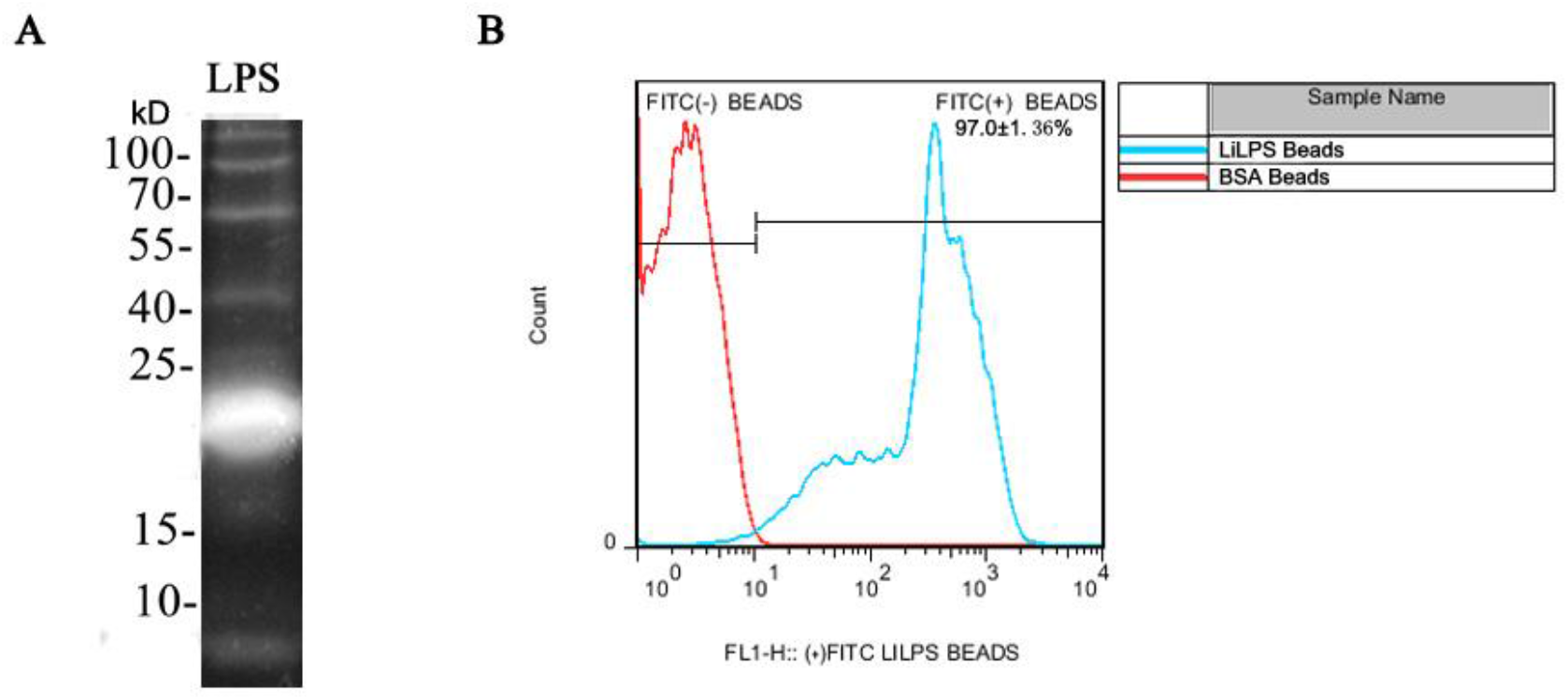
Detection the coating effect of LPS coated beads. **(A)** LPS of *L. interrogans* strain 56606v were extracted by modified phenol water extraction. Pro-Q Emerald 300 was used as fluorochrome to display the LPS band through the reaction of periodate oxidizing saccharide. **(B)** LPS coated beads incubated by *Leptosprira* primary antibody and FITC-conjugated secondary antibody were detected through FCM. Positive rates of coated beads indicating successful coating of specific LPS of *Leptospira* on beads.

**Supplementary Fig. 6.**
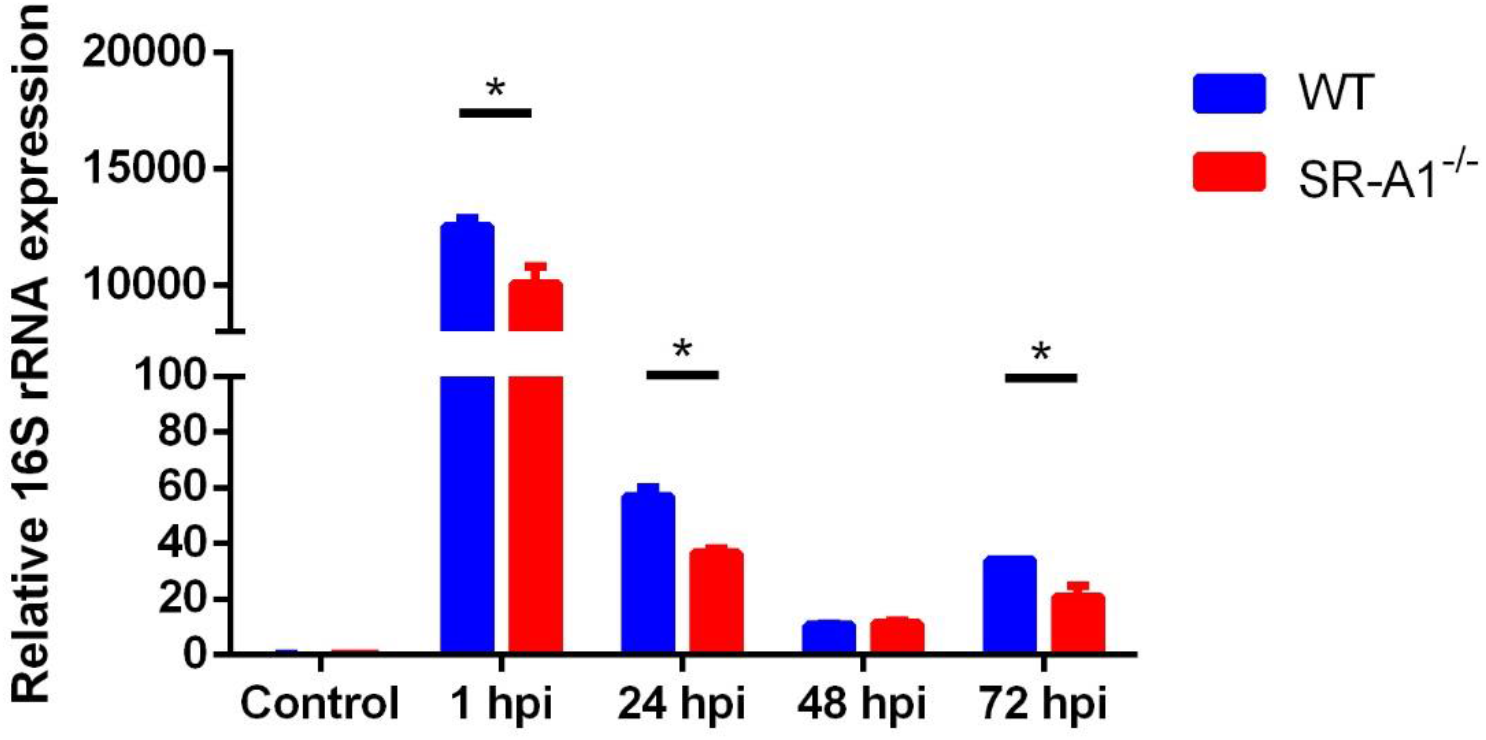
Survival of *L. interrogans* strain 56606v in leptospires infected WT and SR-A1^-/-^ PMs. WT and SR-A1^-/-^ PMs were infected by *L. interrogans* strain 56606v as MOI=100 for 1 h *in vitro*. 100 μg/mL gentamicin was used for 2 h to kill the extracellular leptospires. RNA was extracted from cells of 1 hpi to 72 hpi, and 16S rRNA of leptospires was quantified by real-time PCR. These data were expressed as the mean ± SEM from at least three experiments. *\*P*<*0*.*05*.

**Supplementary Fig. 7.**
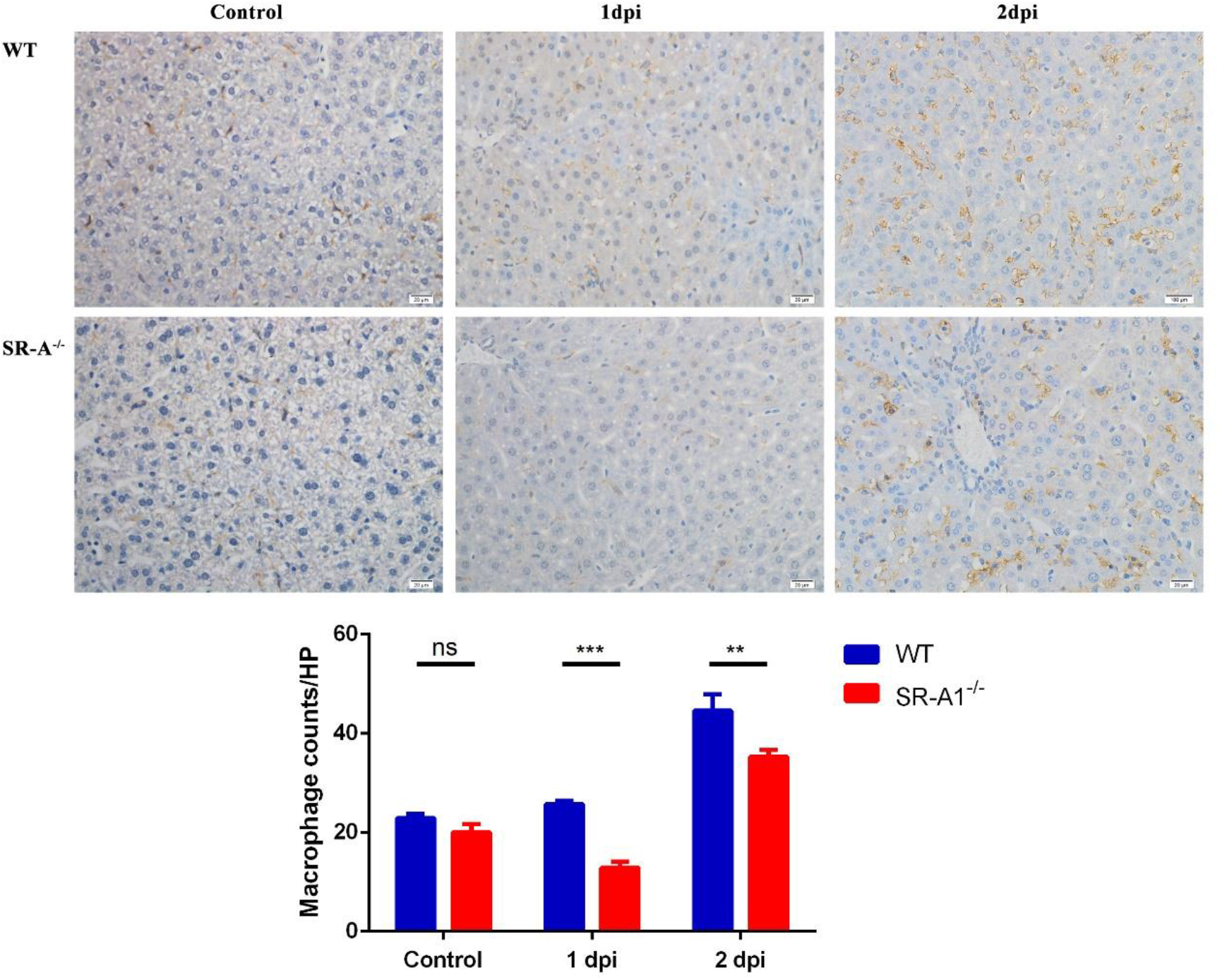
Macrophages recruitment in liver was decreased in SR-A1^-/-^ mice after *Leptospira* stimulation *in vivo*. Livers from WT or SR-A1^-/-^ mice infected with 2×10^8^ leptospires via intraperitoneal route were fixed in neutral buffered 4% formaldehyde, followed by embedding of the tissues in paraffin. Sections were stained for macrophage-specific F4/80 antigen. The average number of F4/80 expressing cells per HP field was calculated. These data were expressed as the mean ± SEM from at least three experiments. *\*\*P*<*0*.*01, ***P*<*0*.*001, ns* no significance.

## Notes

### Competing Interest Statement

The authors have declared no competing interest.

